# IP3 receptor depletion in a spontaneous canine model of Charcot-Marie-Tooth disease 1J with amelogenesis imperfecta

**DOI:** 10.1101/2024.06.03.597092

**Authors:** Marjo K. Hytönen, Julius Rönkkö, Sruthi Hundi, Tarja S. Jokinen, Emilia Suonto, Eeva Teräväinen, Jonas Donner, Rita La Rovere, Geert Bultynck, Emil Ylikallio, Henna Tyynismaa, Hannes Lohi

## Abstract

Inositol 1,4,5-trisphosphate receptors (IP_3_R) mediate Ca^2+^ release from intracellular stores, contributing to complex regulation of numerous physiological responses. The involvement of the three IP_3_R genes (*ITPR1*, *ITPR2* and *ITPR3*) in inherited human diseases has started to shed light on the essential roles of each receptor in different human tissues and cell types. Variants in the *ITPR3* gene, which encodes IP_3_R3, have recently been found to cause demyelinating sensorimotor Charcot-Marie-Tooth neuropathy type 1J (CMT1J). In addition to peripheral neuropathy, immunodeficiency and tooth abnormalities are occasionally present. Here, we report the identification of a homozygous nonsense variant in the *ITPR3* gene in Lancashire Heeler dogs, presenting with a severe developmental enamel defect and reduced nerve conduction velocity. We studied the primary skin fibroblasts of the affected dogs and observed that the nonsense variant in *ITPR3* led to a complete absence of full-length IP_3_R3 protein. Unexpectedly, the protein levels of IP_3_R1 and IP_3_R2 were also markedly decreased, suggesting co-regulation. Functional Ca^2+^ measurements revealed reduced IP_3_R-mediated Ca^2+^ flux upon stimulation of G-protein-coupled-receptors in the affected dog fibroblasts. We were able to rescue the IP_3_R1 and IP_3_R2 depletion by proteasome inhibition but not the IP_3_R3 loss, which was facilitated by nonsense-mediated mRNA decay. These findings highlight the first spontaneous mammalian phenotype caused by a nonsense variant in *ITPR3*, leading to the loss of IP_3_R3. The human and canine IP_3_R3 proteins are highly similar, and our study suggests that the tissue involvement resulting from the receptor’s dysfunction is also conserved. In summary, IP_3_R3 is critical for enamel formation and peripheral nerve maintenance.

**Author summary:** We investigated pet dogs, Lancashire Heelers, with impairments in tooth development and in the nerves that regulate limb muscles. Through genetic studies of the dog pedigree, we found that the phenotypes were caused by a recessively inherited mutation in the *ITPR3* gene, which encodes one of three IP_3_ receptors (IP_3_R) isoforms (IP_3_R3 isoform) that are needed for intracellular Ca^2+^ signaling. Mutated IP_3_R3 has been recently linked to a human inherited neuropathy called Charcot-Marie-Tooth disease type 1J, which impairs peripheral nerve function and is accompanied by immunodeficiency and abnormal teeth in some individuals. We showed that in the skin cells of the affected dogs, the full-length IP_3_R3 protein was completely absent, and also the protein levels of the other two IP_3_R isoforms (IP_3_R1 and IP_3_R2) were severely lowered. This led to impaired agonist-induced Ca^2+^ release and signaling. Our results demonstrate the high conservation between human and canine IP_3_ receptors and their significance for different tissue systems. The genetic studies now highlight that IP_3_R3 is vital for peripheral nerve function and enamel development.

## Introduction

Inositol 1,4,5-trisphosphate receptors (IP_3_Rs) are intracellular Ca^2+^ channels located in the endoplasmic reticulum (ER). Through Ca^2+^ signals induced by IP_3,_ they are involved in a wide range of physiological processes, including differentiation, metabolism, and apoptosis (1, 2). The IP_3_Rs are homo- or heterotetramers composed of three homologs IP_3_R1, IP_3_R2, and IP_3_R3 (encoded by *ITPR1*, *ITPR2* and *ITPR3* respectively), each with different tissue distribution and properties. Studies on IP_3_R knockout animal models and patients with pathogenic variants in IP3Rs have elucidated the importance of distinct isoforms in various tissues (3). Despite their importance, the molecular mechanisms by which IP_3_Rs regulate Ca^2+^ signaling and contribute to various cellular processes are still not fully understood. Nevertheless, the discoveries of disease-associated genetic variants, in addition to genetically modified animal and cell models, support the crucial role of IP_3_Rs (3–9).

The genes encoding IP_3_Rs have been linked to various inherited human diseases. Variants in *ITPR1* cause dominantly inherited spinocerebellar ataxia types 15 and 29, as well as Gillespie syndrome, which exhibits both dominant and recessive inheritance patterns (10–13). A homozygous missense variant in *ITPR2* has been reported to cause isolated anhidrosis (14). *Itpr1* knockout mice either died *in utero* or had severe ataxia and epilepsy after birth (15), whereas *Itpr2* knockouts had reduced sweat secretion (14) and *Itpr3* knockouts had abnormal taste perception (16).

We described dominantly inherited or *de novo ITPR3* variants in individuals with progressive distal muscle weakness, foot deformities and reduced nerve conduction velocity (NCV), a sign of peripheral nerve demyelination (17). Previously, a few other cases of neuropathy associated with *ITPR3* variants of unknown significance had been reported (18, 19). These findings led to a classification of Charcot-Marie-Tooth disease 1J (CMT1J), a new subtype of hereditary neuropathy caused by *ITPR3* variants (Table 1). CMT is a heterogeneous group of inherited peripheral neuropathies affecting the peripheral sensory and motor nerves. The two most common types of CMT are demyelinating CMT1, characterized by reduced NCV (< 38 m/s), and axonal CMT2 (NCV >38 m/s). Recently, two individuals with immunodeficiency were found to carry compound heterozygous variants in *ITPR3*(20). One of those patients had peripheral neuropathy, abnormal tooth eruption and mineralization and thin hair since birth. Furthermore, two unrelated individuals with immunodeficiency from low B and T cells, deciduous teeth, anhidrosis, abnormal gait, bilateral vocal fold paralysis and sensorimotor neuropathy of the lower extremities have been reported (21). Thus, the recent findings suggest that *ITPR3-*related phenotypes extend beyond peripheral nerves.

**Table 1.**
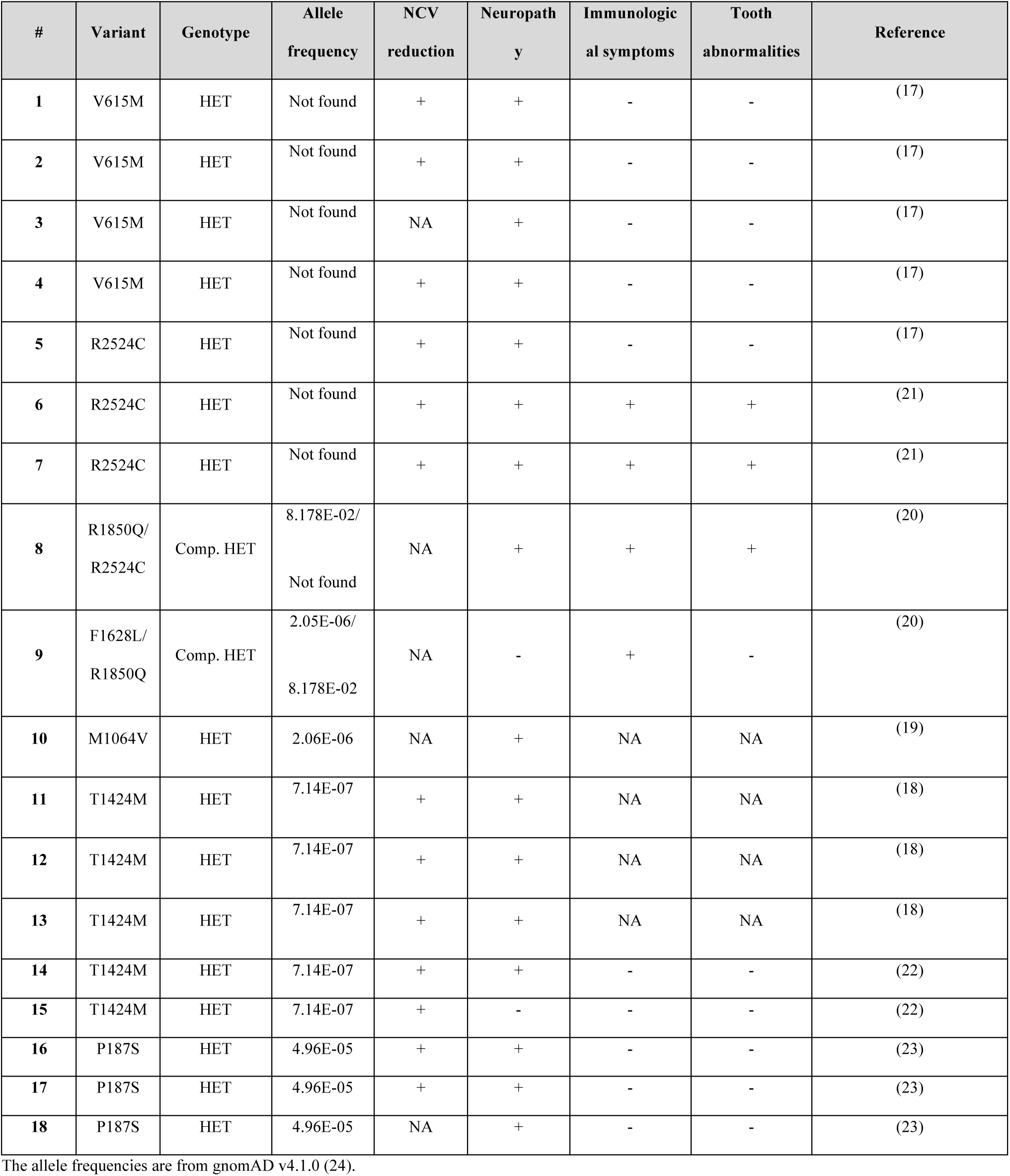
Identified human disease-causing *ITPR3* variants and patient phenotypes.

| # | Variant | Genotype | Allele frequency | NCV reduction | Neuropathy | Immunological symptoms | Tooth abnormalities | Reference |
| --- | --- | --- | --- | --- | --- | --- | --- | --- |
| 1 | V615M | HET | Not found | + | + | - | - | (17) |
| 2 | V615M | HET | Not found | + | + | - | - | (17) |
| 3 | V615M | HET | Not found | NA | + | - | - | (17) |
| 4 | V615M | HET | Not found | + | + | - | - | (17) |
| 5 | R2524C | HET | Not found | + | + | - | - | (17) |
| 6 | R2524C | HET | Not found | + | + | + | + | (21) |
| 7 | R2524C | HET | Not found | + | + | + | + | (21) |
| 8 | R1850Q/<br>R2524C | Comp. HET | 8.178E-02/<br>Not found | NA | + | + | + | (20) |
| 9 | F1628L/<br>R1850Q | Comp. HET | 2.05E-06/<br>8.178E-02 | NA | - | + | - | (20) |
| 10 | M1064V | HET | 2.06E-06 | NA | + | NA | NA | (19) |
| 11 | T1424M | HET | 7.14E-07 | + | + | NA | NA | (18) |
| 12 | T1424M | HET | 7.14E-07 | + | + | NA | NA | (18) |
| 13 | T1424M | HET | 7.14E-07 | + | + | NA | NA | (18) |
| 14 | T1424M | HET | 7.14E-07 | + | + | - | - | (22) |
| 15 | T1424M | HET | 7.14E-07 | + | - | - | - | (22) |
| 16 | P187S | HET | 4.96E-05 | + | + | - | - | (23) |
| 17 | P187S | HET | 4.96E-05 | + | + | - | - | (23) |
| 18 | P187S | HET | 4.96E-05 | NA | + | - | - | (23) |

The allele frequencies are from gnomAD v4.1.0 (24).

In this study, we discovered a homozygous canine *ITPR3* nonsense variant leading to amelogenesis imperfecta and peripheral neuropathy in Lancashire Heeler dogs. This is the first known naturally occurring nonsense variant of *ITPR3* linked to disease, which led to a complete loss of full-length IP_3_R3 protein. Unexpectedly, we found concurrent depletion of IP_3_R1 and IP_3_R2 proteins, which could be rescued by proteasome inhibition. These data point to interdependency in the turnover of the three IP_3_R subunits. Our results suggest a critical role for adequate IP_3_R-dependent Ca^2+^ signaling in enamel development and peripheral nerve function.

## Results

### Enamel defect and peripheral nerve dysfunction in Lancashire Heeler dogs

The affected Lancashire Heeler dogs (Fig 1A) were originally identified by abnormal enamel in the permanent teeth, including severe yellow to brown discoloration, enamel hypoplasia and abrasion leading to exposure of dentin (Fig 1B, C). The enamel defects were already apparent upon tooth eruption. The affected dogs displayed normal breed-typical moving patterns and activity levels. The owners of the affected dogs did not report untypical clumsiness or leg movement in their respective dogs, except for one case at nine years of age. In the general clinical examination, two out of three dogs had deformity detected in their front limbs, i.e. outward rotation of front limbs from the elbow joint. Neurological examination was normal in all affected dogs. No remarkable abnormalities were detected in the complete blood count or serum biochemistry of the examined dogs. Abnormal electromyographic findings consistent with demyelinating neuropathy were detected in all three examined dogs and consisted of fibrillation potentials and positive sharp waves occurring predominantly in distal appendicular limb muscles (i.e. in muscles distal to elbow and knee joints) (Supplementary table 1). Fibrillation potentials were also detected in paraspinal musculature. Motor nerve conduction velocities (MNCVs) were below published canine reference values in both the peroneal and ulnar nerves of the two examined dogs (25). Decreased compound muscle action potential (CMAP) amplitudes and increased CMAP duration (i.e., temporal dispersion) at all stimulation points were observed in these two dogs. One of the examined dogs had normal MNCVs.

**Figure 1.**
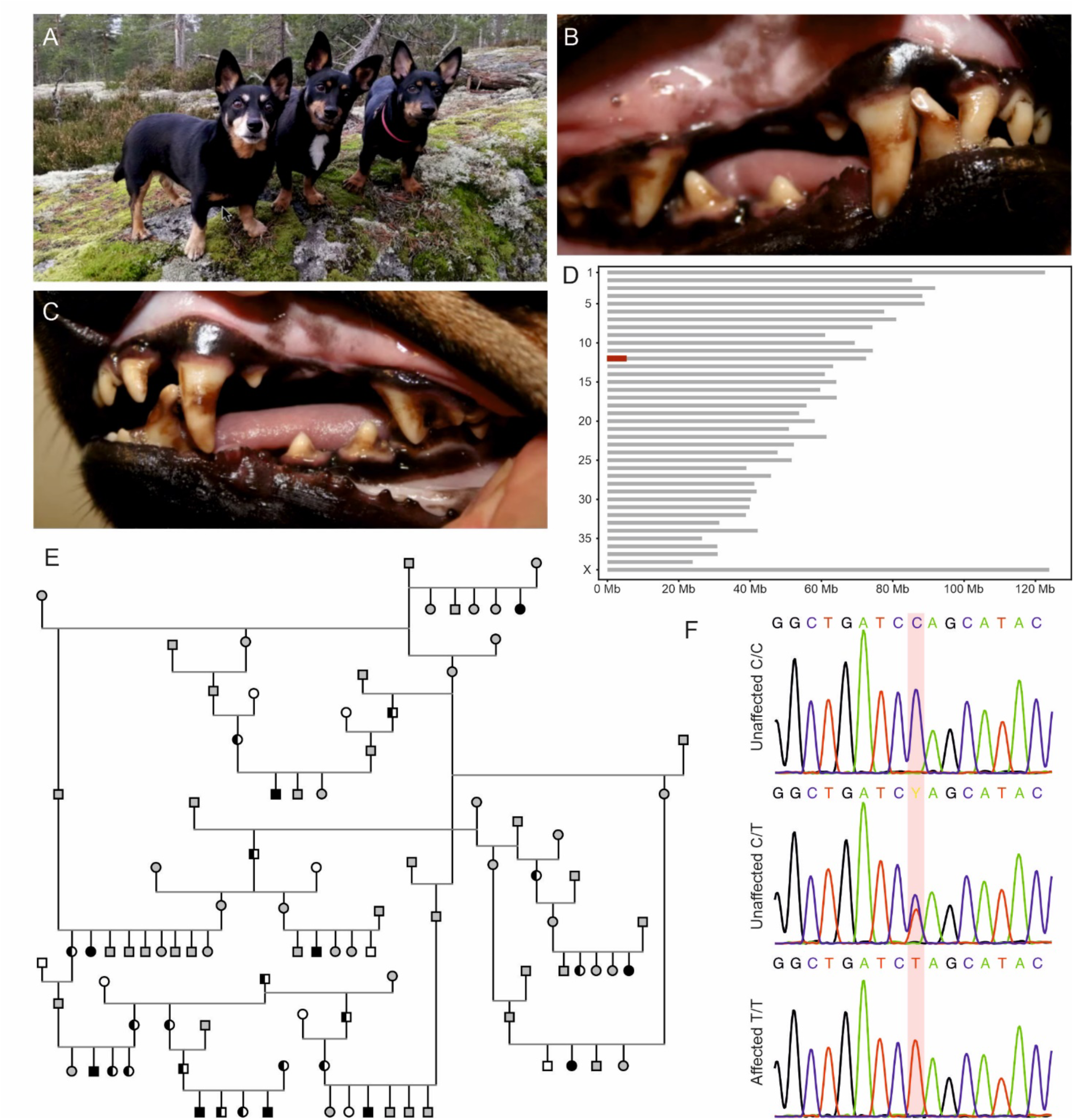
Identification of *ITPR3* p.Q1668X variant in Lancashire Heeler dogs. **(A)** Photograph of Lancashire Heeler dogs, the subject of this study. The image displays the distinct appearance and breed-specific traits of Lancashire Heelers. **(B & C)** Close-up image of an affected Lancashire Heeler dog aged 2 years and 10 months illustrating the typical tooth abnormalities observed in the affected dogs. The typical enamel defects, such as poor enamel formation, discoloration and abrasion of the enamel, are evident. **(D)** Homozygosity mapping of four affected dogs and six controls revealed a case-specific region of allelic homozygosity (ROH) on chromosome 12 (g.156,577-5,127,877) of size 4,97 Mb **(E)** The pedigree compiled around the affected Lancashire Heelers indicates the *ITPR3* variant segregating as a recessive disease. The pedigree includes 10 affected dogs with homozygous (shapes filled with black) *ITPR3* variant, 17 unaffected heterozygous dogs (black and white shapes) and 8 unaffected canines with wildtype *ITPR3* (shapes with black outline). The genotype and phenotype of the rest of the dogs have not been determined (shapes with grey outline). **(F)** Sanger sequencing results confirmed the segregation of the identified *ITPR3* p.Q1668X variant with the phenotype.

### Next generation sequencing revealed a stop-gain variant in *ITPR3*

The pedigree constructed around the affected dogs suggested an autosomal recessive disease (Fig 1E). We carried out a genome-wide genotyping on a cohort of four cases and six controls to identify the runs of homozygosity shared by cases. Homozygosity mapping revealed only one case-specific region of allelic homozygosity (ROH) on chromosome 12 (g.156,577-5,127,877) of size 4,97 Mb (Fig 1D). We performed whole-exome sequencing (WES) on two affected dogs and whole-genome sequencing (WGS) on one affected dog to identify the disease-causing variant. Filtering the SNVs and indels according to a recessive model against variant data of 689 control dogs from other breeds resulted in 4 case-specific homozygous variants (Supplementary table 2). Three of the variants located within the ROH on chromosome 12, however, only one had a predicted effect on the protein: one nucleotide substitution in exon 37 of *ITPR3* (g.12:3214076C>T; XM_038553756.1:5002C>T). The variant substitutes glutamine to a translation termination codon (p.Q1668X) (XP_038409684), which is predicted to result in a significantly truncated protein (amino acids 1668-2690 missing) if translated. The resulting protein would lack part of the coupling region, the transmembrane region and the C-terminal tail, thus lacking the complete C-terminal channel-pore region (26).

We further analyzed the WGS data of the affected dog to exclude any additional non-coding variants, structural variants (SVs), and mobile element insertions (MEIs) using 388 control dogs from other breeds. After filtering the homozygous case-specific variants, 3919 SNVs/indels, 14 MEIs, but no SVs remained (Supplementary Tables 3 and 4). None of the MEIs located within the identified ROH. Of the SNVs and indels, 17 variants were within the mapped ROH, one being the above-mentioned stop-gain variant in *ITPR3.* The remaining variants were non-coding, except for a deletion in ncRNA LOC119874143 annotated by NCBI. However, Ensembl annotated this variant as intronic in *HLA-DRB1*. We investigated the *ITPR3* variant frequency in an additional cohort of 1413 dog genomes sequenced as part of the Dog10K project. All dogs were wild-type, except for one Lancashire Heeler, which was heterozygous for the allele. Based on gene function, predicted pathogenicity and the breed-specific distribution, the nonsense variant in *ITPR3* was prioritized and selected for further validation.

We genotyped the *ITPR3* variant using standard PCR in a cohort of 266 Lancashire Heeler samples, including 10 affected dogs, which confirmed the variant and the segregation of the allele (Fig 1 E, F). The variant was observed as homozygous in all 10 affected dogs and also in one sample from the biobank. According to our records, the additional homozygous dog is deceased and we were unsuccessful in contacting the owner. Therefore, its health status remains unknown. The remaining samples were either heterozygous (n=67) or wild type (n=188). The allele frequency was 16.6% in the study cohort. An additional cohort of over 2.6 million dogs submitted for commercial genetic testing was screened for the *ITPR3* variant to explore the distribution across different breeds. We found only 16 heterozygous dogs, 13 of which had undergone DNA-based breed ancestry testing. Ten of these heterozygotes were purebred Lancashire Heelers, two were mixed breed dogs with detected genomic ancestry from the Lancashire Heeler (30% and 2% contributions, respectively), and one was a dog of highly mixed ancestry.

The IP_3_R1, IP_3_R2, and IP_3_R3 are highly similar proteins between humans and canine, with 96%-98% of the amino acids being identical between the species (Fig 2A) (Supplementary Fig 1). The premature stop codon caused by the variant resides in a conserved site in the IP_3_R3 regulatory/coupling region upstream of the channel region containing the channel pore (Fig 2B). The possible truncated protein would be non-functional without transmembrane alpha helices or the channel pore (Fig 2C).

**Figure 2.**
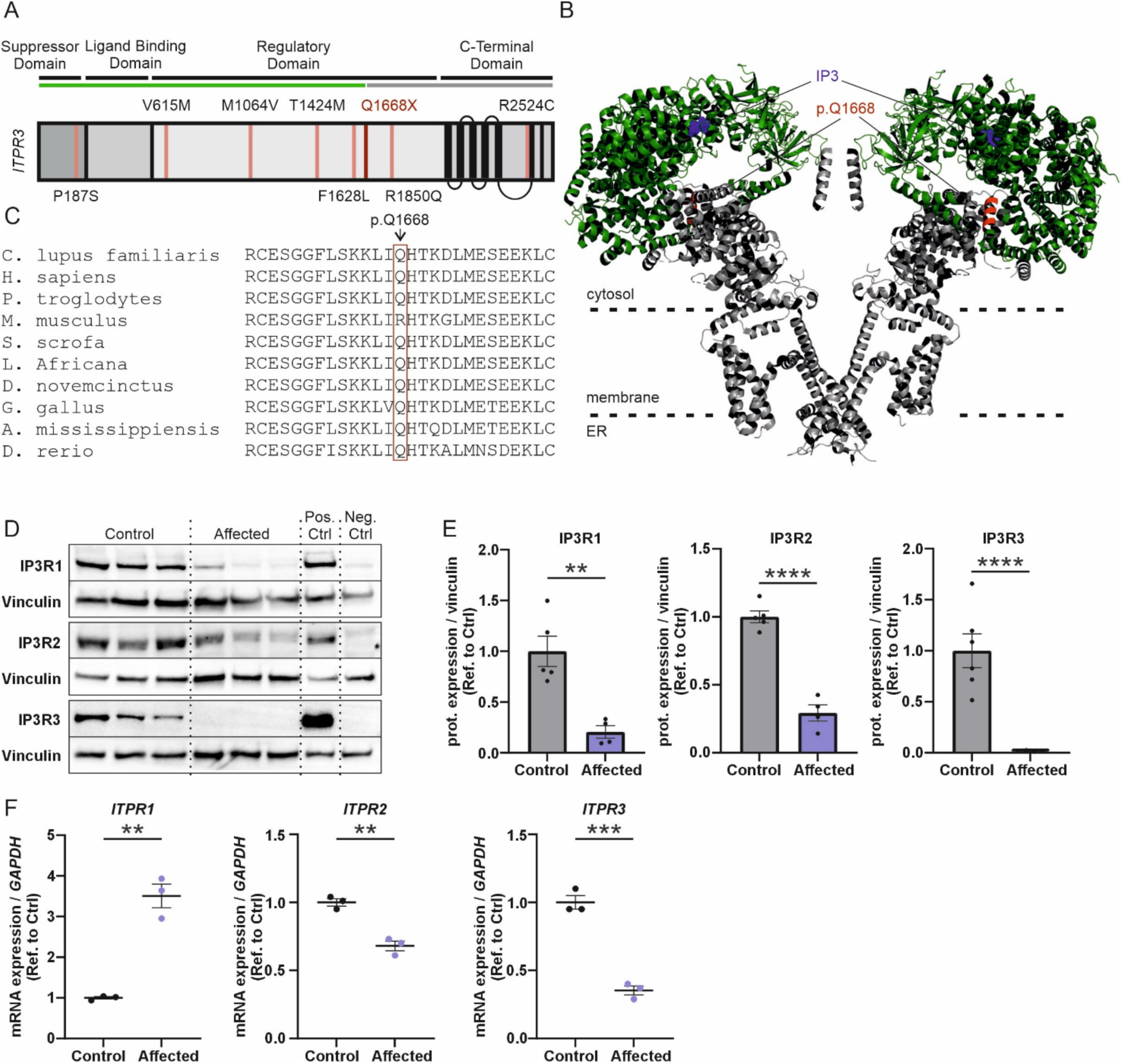
The affected dogs have a complete loss of IP_3_R3 and reduced IP_3_R1 and IP_3_R2 protein levels (A) Schematic representation of the secondary structure of IP_3_R3 protein. The functional domains are annotated, highlighting the key functional domains of the protein. These domains include N-terminal suppressor domain, IP_3_-binding ligand binding domain, regulatory domain, and C-terminal domain. Known human CMT1J variants are depicted at their respective locations in IP_3_R3 in black, and the dog variant in red. The length of the potential truncated protein is depicted in green and non-translated region in grey. (B) The position of the p.Q1668X variant in IP_3_R3 3D structure. The canine p.Q1668X corresponds to the human p.Q1649. The variant site and five adjacent amino acids are highlighted in red. Two IP_3_R3 subunits are illustrated. The potential truncated protein is depicted in green and the non-translated region in grey. IP_3_ in the binding site is depicted in blue. Human IP3R3 3D model: (7T3T) (52). (C) The canine p.Q1668X variant site and amino acid conservation. (D) Representative Western blots showing the reduced IP_3_R1 and IP_3_R2 protein levels and loss of IP_3_R3 in the affected dogs. (E) Quantification of the Western blot analysis of the IP_3_R1-3 protein levels. Mean ± SEM, n= 4-6 replicates. (F) Quantification of RT-qPCR analysis of the *ITPR1*, *ITPR2* and *ITPR3* mRNA expression levels. Mean ± SEM, n = 3 replicates. Statistics are analyzed by Student’s t-test (*p < 0.05, **p < 0.01, ***p < 0.001).

### Complete loss of IP_3_R3 and reduced IP_3_R1 and IP_3_R2 levels in fibroblasts of affected dogs

To investigate if the p.Q1668X variant leads to truncation of IP_3_R3, we obtained primary skin fibroblasts from the affected Lancashire Heelers and control dogs and performed a Western blot with an antibody directed against the N-terminal (22-230 amino acid residue) region of IP_3_R3 (Fig 2D, E). The affected dog fibroblasts showed a complete lack of full-length IP_3_R3 proteins. In addition, no N-terminal fragment was visible around 170-180 kDa, the theoretical molecular weight of the truncated protein fragment (Supplementary figure 2), suggesting that the nonsense-mutant *ITPR3* mRNA is degraded by nonsense-mediated decay. Indeed, RT-qPCR analysis showed a reduction of *ITPR3* mRNA level (reduced to 35% of control level) in the affected dog fibroblasts (Fig 2F). Surprisingly, we also found decreased protein levels of IP_3_R1 and IP_3_R2 in the affected dog fibroblasts (79% and 71% reduction, respectively). Also, the mRNA level of *ITPR2* was decreased, whereas the level of *ITPR1* mRNA was increased in the affected dog fibroblasts.

These results were consistent with partial nonsense-mediated mRNA decay caused by the p.Q1668X variant in *ITPR3*, and if any truncated protein was produced, it was not stable. Elevation of *ITPR1* mRNA may represent a compensatory upregulation of the alternative IP_3_R, which ultimately appeared futile as the protein levels of both IP_3_R1 and IP_3_R2 were decreased in the affected Lancashire Heeler dog fibroblasts. The latter result was unexpected and prompted further investigation of IP_3_R turnover.

### Proteasome inhibitor MG-132 rescues the IP_3_R1 and IP_3_R2-protein levels

To assess the mechanism of IP_3_R depletion, we treated affected Lancashire Heeler fibroblasts with proteasome inhibitor MG-132 for 12 and 24 hours. After the MG-132 treatment, we analyzed the protein levels of IP_3_R1-3 using Western blotting with IP_3_R-isoform-specific antibodies (Fig 3A). The analysis revealed significant changes in the IP_3_R1 and IP_3_R2 levels compared to non-treated cells (Fig 3B). The MG-132 treatment rescued the depletion of IP_3_R1 and IP_3_R2 to levels comparable to those in the control cell lines, while the treatment had no consistent effect on the protein levels in the control cells. As anticipated, the levels of the full-length IP_3_R3 protein were not affected by MG-132. Hence, the depletion of IP_3_R1 and IP_3_R2 proteins appears to be caused by proteasomal degradation, as MG-132 restored their protein levels.

**Figure 3.**
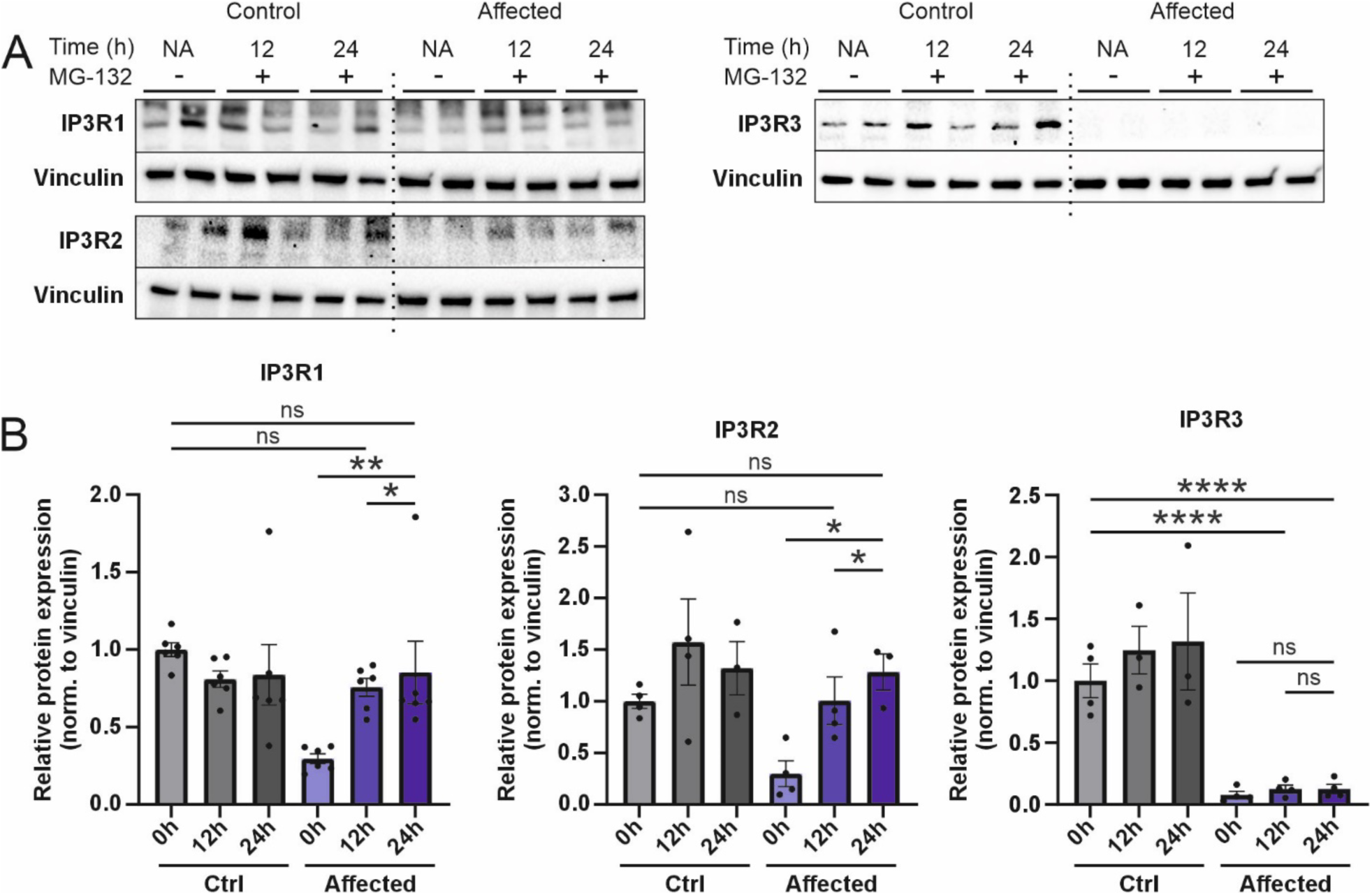
Proteasome inhibitor MG-132 rescues the IP_3_R1 and IP_3_R2 protein levels **(A)** Representative Western blots of the IP_3_R1-3 proteins after treating the cells with the proteasome inhibitor MG-132 for 12 and 24 hours. **(B)** Quantification of the Western blot analysis of the IP_3_R1-3 expression levels after MG-132 treatments. n = 3-6 replicates. Data are presented as mean ± SEM. Statistics are analyzed by one-way ANOVA with Dunnett’s multiple comparison test (*p < 0.05, **p < 0.01, ***p < 0.001).

### The p.Q1668X variant leads to a reduction in IP_3_R-mediated cytosolic Ca^2+^ release

To assess the effect of the *ITPR3* nonsense variant on Ca^2+^ signaling, we utilized functional Ca^2+^ imaging on the affected Lancashire Heeler dog fibroblasts and control cell lines. We analyzed the dog fibroblasts with ratiometric Fura-2 AM Ca^2+^ indicator using a fluorescent microscope. The measurements were performed in the absence of extracellular Ca^2+^ to ensure that Ca^2+^ signals originated from intracellular stores. We measured the cytosolic [Ca^2+^] rises in living cells exposed to the G-protein-coupled-receptor (GPCR) agonist ATP, while the total intracellular releasable Ca^2+^ was estimated using the Ca^2+^ ionophore ionomycin.

First, we elicited the IP_3_ production of the cells by 80 µM ATP and measured the cells’ capacity to generate cytosolic Ca^2+^ signals upon production of IP_3_. The affected Lancashire Heeler fibroblasts had significantly reduced ATP-evoked Ca^2+^ responses compared to the control cells, as both the maximum amplitude and area-under-the-curve analysis displayed similar decreases in the affected fibroblasts (Fig 4A). We then assessed the total intracellular Ca^2+^ storage with 2 µM Ca^2+^ ionophore ionomycin. There were no differences between the affected and the control fibroblasts, indicating comparable intracellular Ca^2+^ loading between the cell lines (Fig 4B).

**Figure 4.**
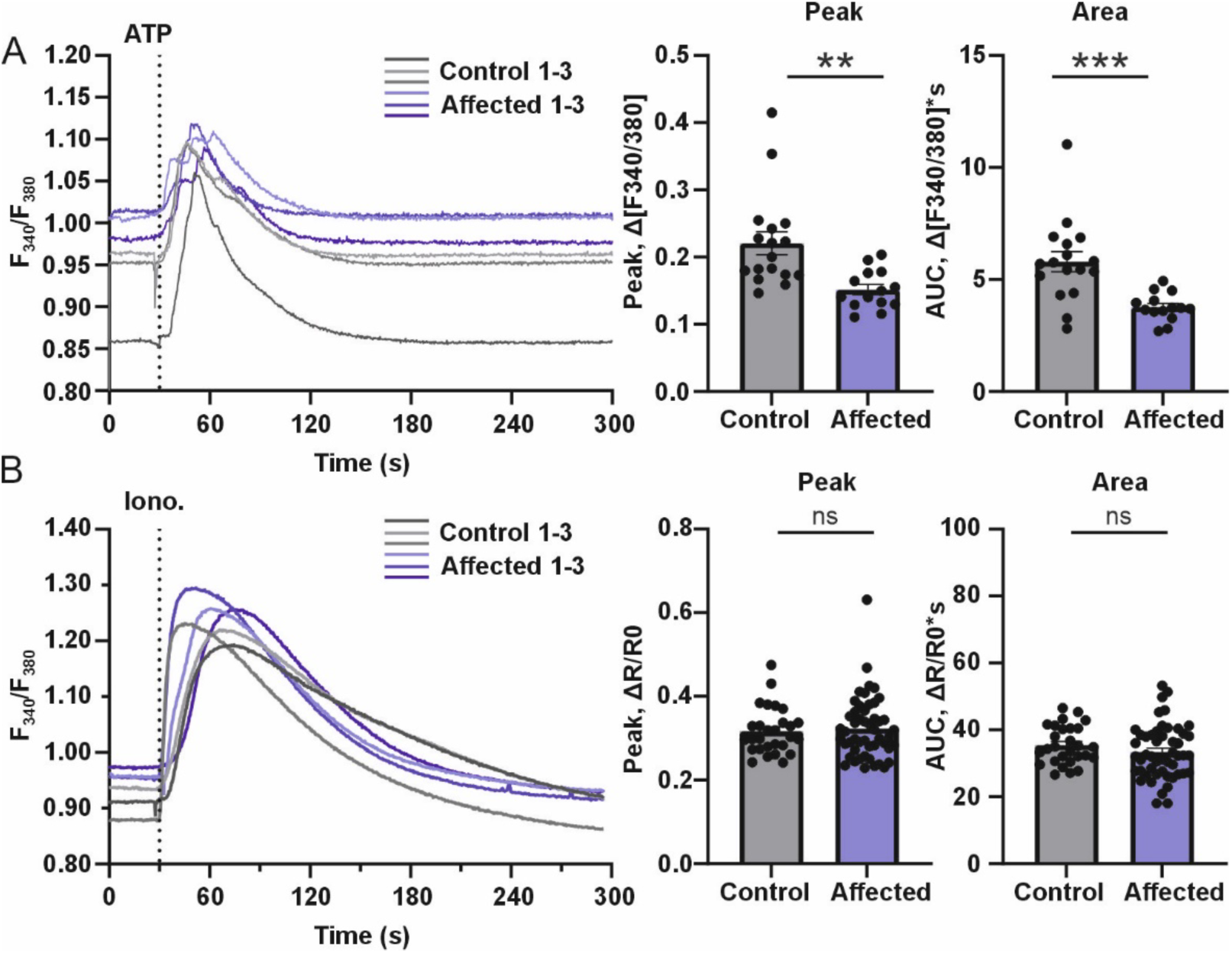
The Q1668X variant leads to reduction in IP_3_R-mediated Ca^2+^ release. The Ca^2+^ flux measurements were performed with ratiometric fluorescent Ca^2+^ indicator FURA-2 AM. The Ca^2+^ flux analysis was performed with one homozygous Lancashire Heeler dog skin fibroblast line and one control dog cell line. **(A)** Fluorescent response curves of the affected and control cell lines after stimulation with GPCR agonist ATP. Average fluorescent response curves of the affected and control cell lines after stimulation with 80 µM ATP. The affected fibroblast cells had significantly reduced peak amplitude and AUC (n = 36 - 47 cells, 3 independent experiments). **(B)** Fluorescent response curves of the affected and control cell lines after stimulation 2.5 µM ionomycin. There were no significant differences in the peak amplitude or AUC between the affected and control cells (n = 73 – 118 cells, 3 independent experiments). Data are presented as mean ± SEM. Statistics are analyzed by Student’s t-test (*p < 0.05, **p < 0.01, ***p < 0.001).

Next, we analyzed the dog fibroblasts with Ca^2+^ indicator Cal-520 AM using the drug screening system FDSS/µCell. The Lancashire Heeler fibroblasts had a significantly reduced response to 100 µM ATP, similarly as observed with ratio metric measurements (Supplementary figure 3A). We then assessed the total ER Ca^2+^ storages with 2 µM irreversible SERCA inhibitor thapsigargin and the total intracellular Ca^2+^ stores with 2.5 µM ionomycin. These results indicate that the Ca^2+^ loading of intracellular Ca^2+^ stores and ER was significantly different between the affected and control fibroblast cells (Supplementary figure 3B & C).

Altogether, these results indicate that the p.Q1668X variant caused a loss of IP_3_R3 and affected the IP_3_R1- and IP_3_R2-protein levels through proteasomal degradation, resulting in deranged Ca^2+^ signaling.

## DISCUSSION

We report the first spontaneous large animal model of human CMT1J with a nonsense mutation in *ITPR3*, leading to a complete loss of full-length IP_3_R3 proteins in Lancashire Heeler dogs. The results provide additional evidence for the critical role of IP_3_R3 in the development and function of the peripheral nerves and teeth.

IP_3_Rs are major regulators of intracellular calcium signaling, however, the functional importance of each of the three receptors in different tissues and cell types is not fully established. Earlier studies of knockout mice have yielded important insights into the role of the different IP_3_R isoforms in physiology, but whether such findings translate to humans has not been known. Genetic studies identifying pathogenic variants in the IP_3_R genes have now elucidated that each human IP_3_R has unique tissue-specific roles that the other two receptors cannot compensate for. IP_3_R1 has a significant role in the central nervous system, leading to ataxia phenotypes when mutated (3), whereas IP_3_R2 is critical in sweat glands, causative of anhidrosis and heat intolerance (14). IP_3_R3, on the other hand, has recently been linked to peripheral neuropathy, with some patients having additional immunodeficiency and tooth abnormalities (17, 20, 21).

Here, we describe the first spontaneous model of human CMT1J with significant clinical similarities. The affected dogs showed no systematic clinical evidence of motor or sensory impairment, but the electrodiagnostic examinations revealed neurogenic changes in muscle and decreased NCV. These findings were consistent with subclinical peripheral neuropathy. The human disease CMT1J, caused by pathogenic variants in *ITPR3*, is typically a slowly progressive sensorimotor neuropathy (17–19). Electrodiagnostic studies in the human patients have shown decreased NCV, consistent with demyelination, which has also been confirmed in sural nerve biopsy (17). The demyelinating CMT type suggests the involvement of Schwann cells, which could indicate that the role of IP_3_R3 in peripheral nerve maintenance is through a defect in the myelinating cells. It is noteworthy that the

severity of CMT1J varies between patients. Analogously to the dogs described here, one individual in her 50s who carried the *ITPR3* p.T1424M variant had no signs of neuropathy but widespread slowing of NCV, confirming the presence of subclinical neuropathy (22).

The affected Lancashire Heelers had features of amelogenesis imperfecta, including significant discoloration of teeth, enamel hypoplasia and abrasion leading to dentin exposure. Similarly, signs of tooth abnormalities have been reported in some human patients with the *ITPR3* p.R2524C variant (20, 21). Ca^2+^ is a critical component in enamel mineralization, as it is essential for the formation and growth of hydroxyapatite crystals, which provide the structural integrity and hardness of enamel in teeth (27, 28). Ca^2+^ signaling pathways are involved in the differentiation and maturation of ameloblasts, the cells responsible for enamel formation (29, 30). IP_3_Rs are key Ca^2+^ regulators in ameloblasts, functioning as ER’s main Ca^2+^ release channel (31).

Other important Ca^2+^ handling proteins involved in ameloblast Ca^2+^ signaling are ORAI1 and STIM1, responsible for Ca^2+^ uptake and refilling ER Ca^2+^ storages via store operated calcium entry (SOCE) (32). Loss-of-function variants in *STIM1* and *ORAI1* cause calcium release-activated channelopathies (CRAC) with ectodermal dysplasia, leading to enamel defects and anhidrosis, immunodeficiency, and muscular hypotonia with muscle weakness (33). The complex interplay between IP_3_Rs, ORAI1, and STIM1 in amelogenesis and dental enamel development explains the similarities between their genetic defects. However, our results of the *ITPR3* nonsense mutation in Lancashire Heeler dogs, leading to the absence of IP_3_R3, suggest that it is the most important IP_3_R for enamel formation, which is also supported by the lack of tooth phenotypes in human *ITPR1* or *ITPR2* diseases.

Unlikely in the studied dogs, CMT1J is predominantly an autosomal dominant disease in humans, but a few patients with compound heterozygous *ITPR3* mutations have been reported (20). The human p.V615M and p.T1424M variants seem to lead to a purely neuropathic phenotype. The differences in patient phenotypes might stem from the severity of each variant in the channel function, as suggested by the results of a study where the mutant proteins were expressed in cells lacking all wild-type IP_3_Rs (34). The above-mentioned neuropathy-causing mutants formed abnormal but partially functional IP_3_R channels. In contrast, the p.R2524C variant locating in the channel pore led to a complete absence of channel function upon stimulation (34). We showed that the *ITPR3* nonsense variant caused a loss of IP_3_R3 in dog fibroblasts. It may thus be functionally similar to the dead-pore human p.R2524C variant, although the cells harboring p.R2524C can still form IP_3_R tetramers and localize into the ER membrane. Tetramers containing the dead-pore variant may have lost Ca^2+^ conduction but could produce additional harmful signaling effects, which are not seen in dogs. The mutation type may also contribute to the lack of immunological phenotype in the affected dogs, which has been reported in a few human patients (20, 21). Together, these data suggest a genotype-phenotype relationship for *ITPR3* variants in humans and dogs, where the mildest dominantly inherited variants may lead to neuropathy while more severe variants produce multisystem involvement.

We unexpectedly observed that besides the loss of IP_3_R3 in the affected dogs’ fibroblasts, the protein levels of IP_3_R1 and IP_3_R2 were also severely reduced. The contribution of these additional deficiencies in IP_3_Rs to the dog phenotype is not known. Furthermore, we only had access to skin biopsies of the pet dogs and cannot rule out if IP_3_R1 and IP_3_R2 were less or more affected in other cell types of the affected dogs. The loss of IP_3_R3 was most likely mediated at least partially by nonsense-mediated mRNA decay, and if any truncated protein was produced, it could not be detected by Western blotting. What could explain the reduction in IP_3_R1 and IP_3_R2 levels when the production of IP_3_R3 was impaired? IP_3_Rs are known to have a short half-life once activated, which is achieved through their rapid degradation by the ER-associated degradation (ERAD) pathway, which is ubiquitin-proteasome dependent (35–37). This mechanism may shield cells from the deleterious effects of overactivation of Ca^2+^ signaling pathways. One possibility is that the truncated IP_3_R3 may be a misfolded ER-protein that constitutively activates the ERAD pathway, and the overactivation of the system accelerated the degradation of IP_3_R1 and IP_3_R2. The observed rescue by the proteasomal inhibitor could support this hypothesis. Finally, the reduction in ATP-evoked Ca^2+^ responses was moderate in the skin fibroblasts of the affected dogs, suggesting that the residual IP_3_Rs were functional on a level that was likely sufficient in most tissues.

In conclusion, we have described the first nonsense mutation in *ITPR3,* causing subclinical neuropathy and abnormal enamel development in Lancashire Heeler dogs. This study extends the phenotypic range of defective IP_3_R3 function due to disease-linked variants, bridging findings emerging from recent genetic studies in human patients. Our results point to the role of IP_3_R3 in enamel formation and link *ITPR3* variants to CRAC channelopathies.

## METHODS

### Experimental model and subject details

#### Cell lines and culture conditions

All dogs in this study were privately owned pet dogs. The dog owners gave written informed consent to perform the experiments. The skin biopsies from two affected and four unaffected dogs were collected for fibroblast culture. Skin biopsies were performed under general anesthesia with a disposable sterilized biopsy dermal anchor puncher. The fibroblasts were cultured in Dulbecco’s Modified Eagle Medium (DMEM), supplemented with 10% FBS (Life Technologies), 1% penicillin/streptomycin (Life Technologies), 1% L-glutamine (Life Technologies) and 0.2% uridine (Sigma). Cells were incubated at 37 °C in 5 % CO_2_.

### Method details

#### Study cohort

EDTA-blood samples were collected from 266 Lancashire Heeler dogs, including 10 affected dogs, their relatives, and other dogs in the canine biobank at the University of Helsinki. The samples were stored at -20°C until genomic DNA was extracted using the semi-automated Chemagen extraction robot (PerkinElmer Chemagen Technologie GmbH). DNA concentration was determined by either DeNovix DS-11 Spectrophotometer (DeNovix Inc., Wilmington, Delaware, USA) or Qubit 3.0 Fluorometer (Thermo Fisher Scientific Inc.). All sampled animals were privately owned pet dogs enrolled in the study with the owner’s informed consent. Sample collection was ethically approved by the Animal Ethics Committee of State Provincial Office of Southern Finland (ESAVI/343/04.10.07/2016 and ESAVI/25696/2020). The pedigree was compiled using GenoPro genealogy software.

#### Clinical examination

Three dogs underwent a physical and neurologic examination and a bloodwork assessment, including complete blood count and serum biochemistry. The neurological examination included an evaluation of the dogs’ mental status, posture, gait, an assessment of postural reactions, spinal reflexes, panniculus reflex, perineal and bulbouretral reflex, and an evaluation of cranial nerves.

The dogs were anesthetized for electrodiagnostic examination. We performed all electrophysiological examinations with a Cadwell electrodiagnostic machine (Cadwell Industries, Inc., Kennewick, WA). EMG was performed bilaterally in the proximal and distal muscles of the pelvic and thoracic limbs and the paraspinal muscles. A disposable concentric needle electrode was used for EMG analysis and a subdermal needle electrode placed subcutaneously on the animaĺs flank served as the ground. Abnormalities detected included abnormal insertional and spontaneous activity, such as fibrillation potentials and positive sharp waves.

MNCV of ulnar and peroneal nerves were measured unilaterally performed with two stimulating monopolar needle electrodes. Subdermal needle electrodes served as a recording electrode, a reference electrode, and a ground electrode. The rectal temperature of the dog was measured during the electrodiagnostic testing.

#### Homozygosity mapping

We genotyped four affected and six unaffected dogs using Illumina’s CanineHD Whole-Genome Genotyping BeadChip containing 173,662 markers (Lincoln, NE, USA). We conducted the pre-analytical QC using PLINK (version 1.96b6.20) (38) and included pruning for a sample call rate of > 95 %, marker call rate of > 95 %, and Hardy-Weinberg equilibrium p-value < 1×10^-8^. Pruning for minor allele frequency or LD was not performed as recommended by Meyermans et al. (39). After QC, all four cases and six controls, as well as 165,826 markers, remained for analysis.

Detection of runs of homozygosity (ROH) was performed with PLINK 1.9 (38). We optimized the population-dependent parameters using simulated data, and then detected ROH shared by the affected dogs. The minimum marker size for ROH (--homozyg-snp) was calculated based on the formula described by Purfield et al. (40) with α = 0.05, N_s_ = 149084, N_i_ = 6 and mean SNP heterozygosity = 0.27. The parameters --homozyg-window-snp, --homozyg-window-missing, --homozyg-window-het, --homozyg-window-threshold and --homozyg-kb were set to a fixed value depending on the analysis (Supplementary table 5).

To establish suitable values for ROH density and maximum gap, genotypes for a fully homozygous individual were simulated based on the population map file and analyzed for maximal genome coverage as described by Meyermans et al. (39). To calculate maximum genome coverage for the simulated genome, --homozyg-gap was set to 2000 kb and --homosyg-density to 200 kb/snp; genome coverage was determined as the total length of the resulting ROH. The simulated genome was then analyzed by varying --homozyg-density from 10 to 125 kb/snp in increments of 5 kb and --homozyg-gap from 20 to 1000 kb in increments of 20 kb (Supplementary table 5).

Based on the simulation, density was set to 25 kb/snp and maximum gap to 200 kb, as genome coverage reached 100 % for --homozyg-density at 30 and increased only negligibly for --homozyg- gap after 200. The detection of ROH was performed with --homozyg-group using these parameters. The ROH included in the further analyses were required to be allelically shared by all four affected dogs and either absent or allelically different in the control dogs.

#### Whole-genome and exome sequencing

We performed exome sequencing on two samples, where the exome library was prepared with 140702_canFam3_exomeplus_BB_EZ_HX1 kit. The prepared library, with a total capture size of exome of 152 Mb obtained from the Roche NimbleGen SeqCap EZ target enrichment design (41) was sequenced on Illumina NextSeq500 platform at Biomedicum Functional Genomics Unit. The exome sequence fastq, with a read length of 300 bp, was aligned to GSD1.0 (42) with Y chromosome appended from ROS_Cfam_1.0 using bwa-mem and had 49X and 52X coverage. Coverage was calculated by the percentage of mapped reads on the target captured region. Duplicates were marked using Mark duplicates feature in GATK and Indel realignment was performed on the aligned bam files, followed by base quality score recalibration and variant calling in gVCF mode using GATK Haplotype caller (43). Whole-genome sequencing (WGS) was conducted on another affected dog on Illumina HiSeq2000 high-throughput sequencing platform with a read length of 300 bp and an average coverage of 19X. Post quality control using fastqc, the reads from the WGS sample were processed using SpeedSeq open-source software with bwa (v0.7.15) for alignment. Similar to the exome processing, duplicates were marked, followed by indel realignment and variant calling in gVCF mode. The gVCFs were combined using combineGVCFs and jointgenotyped using GenotypeGVCFS in GATK version 4.2. The combined VCFs files were annotated for genomic and functional annotations using annovar (44) with Ensembl release100 and NCBI *Canis lupus familiaris* Annotation Release 106. For the WGS sample, mobile element insertions were detected using Mobile Element Locator Tool (MELT) (45). Manta 1.6 (46) was used to identify Structural Variants (SVs), which included insertions, deletions, inversions, and duplications. The identified SVs were genotyped using Graphtyper2.

#### Variant analysis

The detected variants were imported into an in-house server with an adapted version of webGQT for canine variants for inheritance model-based variant filtering to identify the casual variant (47). We performed the analysis of the variant data separately for SNVs and indels and, SVs and MEIs. We filtered SNV and indel variant data of three cases (two dogs with WES and one with WGS data) against 689 control genomes assuming an autosomal recessive inheritance. Specifically, the affected dogs were assumed to share the variants in a homozygous state, while no heterozygous or homozygous calls were allowed per alternative allele in the controls. Whole-exome sequencing and WGS data from 689 dogs from different breeds and 4 wolves with 25X average depth (17-59X) were utilized for filtering the SNVs and indels, while 388 of these samples were available for MEI and SV analysis. Additionally, we used 1413 control genomes that had been sequenced (20X average depth) as a part of the Dog 10K genomes project (48). The variant data of all control genomes were aligned and called as described above.

NCBI transcript XM_038553756.1 and protein XP_038409684.1 were used to count the nucleotide and amino acid positions for *ITPR3*, respectively. The protein alignment was performed with the Clustal Omega algorithm (https://www.ebi.ac.uk/Tools/msa/clustalo).

#### Variant validation

Genotyping of individual dogs was performed with PCR, followed by Sanger sequencing. The DNA template was amplified with Taq polymerase (Biotools B&M Labs, S.A.) The primers used are listed in Supplementary Table 6. The products were treated with exonuclease I (New England Biolabs) and rapid alkaline phosphatase (Roche Diagnostics) and then sequenced using the PCR primers on an ABI 3730 capillary sequencer (Life Technologies). The sequence data were analyzed using Unipro UGENE v1.32.0 (49–51)

A sample of 2,651,699 pure-bred and mixed breed dogs, collected via owner submission for commercial testing, was screened using the Wisdom Panel^TM^ (Wisdom Panel, WA, USA) genetic testing platform, including breed detection assessment.

#### Western blot

Skin fibroblasts were cultured until they reached 70-80% confluency and lysed after collection with RIPA buffer (Cell Signaling #9806). Aliquot containing 20 µg of total protein was boiled at 95 °C, separated in 4-20% Criterion™TGX™ gels (Bio-Rad), and transferred to 0.2 µm nitrocellulose (Bio-Rad) using Trans-Blot Turbo (Bio-Rad). The membranes were blocked in 10% milk in 0.1% TBS-T. Antibodies used were: IP_3_R1^(CT-1)^, IP_3_R2^(NT-2)^ (gift from Dr. David Yule, University of Rochester, NY), IP_3_R3 (BD-Transduction #610312) and Vinculin (Sigma #V9264-25UL). Secondary antibodies were: Anti-rabbit and anti-mouse (Jackson Immunoresearch#111-035-144 and #115-035-146). The blots were imaged with WesternBright™ ECL-spray (Thermo Scientific) and Molecular Imager ChemiDoc XRS+ (Bio-Rad).

#### Real-time quantitative PCR

The RNA was extracted from fibroblast by NucleoSpin® RNA extraction kit (Macherey-Nagel #740955) and reverse transcribed by Maxima first strand cDNA synthesis kit (Thermo Fischer) using Bio-Rad CFX Maestro 1.1 software (Bio-Rad). The primers used are listed in Supplementary table 6.

#### Functional Ca^2+^ imaging

We investigated IP3R-mediated Ca^2+^ signaling using non-ratiometric manual Ca^2+^ assay with Ca^2+^ dye Cal-520 AM and ratiometric automated Ca^2+^ assay with Ca^2+^ dye FURA-2 AM. The non-ratiometric Ca^2+^ signaling dynamics of the affected Lancashire Heeler fibroblast cells and the control cells were analyzed using the FDSS/µCell functional drug screening system (Hamamatsu). The fibroblasts were seeded in 96-well plates (Greiner Bio-one #655090) and analyzed using Ca^2+^ indicator dye Cal-520 AM (Aat Bioquest). The cells were loaded with 1 µM Cal-520 and incubated at 37 °C for 30 min in modified Krebs solution (150 NaCl, 5.9 KCl, 1.2 MgCl_2_, 11.6 HEPES (pH 7.3), 1.5 CaCl_2_, in mM). After incubation, the cells were washed once with the buffer and incubated for 30 min at 37 °C to allow dye de-esterification. After the second incubation, the cells were washed once with the buffer and imaged at 37 °C with Krebs solution with 0 mM Ca^2+^. The 3 mM EGTA was added to the cells after 30 seconds, and after an additional 60 seconds, the cells’ Ca^2+^ response was induced by either 100 µM ATP, 2 µM thapsigargin or 2.5 µM ionomycin, and imaged for 540 seconds. In experiments with ATP and thapsigargin, the cells were additionally exposed to ionomycin after 390 seconds. The Ca^2+^ signaling experiments were quantified with GraphPad Prism 9 software. The fluorescent signal was then normalized to the baseline values, assessed by the fluorescent signal 60 sec prior to the stimuli.

For the ratiometric Ca^2+^ assays, the dog fibroblast cells were seeded in MatTek glass bottom dish (MatTek #P35G-1.5-14-C) and imaged with Zeiss Axio Observer Z1 inverted phase contrast fluorescence microscope. The cells were loaded with 1 µM Fura-2 AM (ThermoFisher) in a fibroblast culture medium at 37 °C for 30 min. After loading, the cells were washed twice with Krebs solution and let de-esterificate for 15 min before imaging at 37 °C. The cells were washed twice before imaging with Ca^2+^-free Krebs solution containing 3 mM EGTA and stimulated with either 80 µM ATP or 2 µM ionomycin after 30 seconds of imaging. The cells were masked, and the mean pixel intensity was measured for each frame using FIJI ImageJ software. The baseline values were selected in the first 30 seconds, and the relative intensities were calculated. The results were analyzed using Clampfit software (Axon instruments). The AUC of the peak and the peak amplitude were measured and the cells with no response (<10% increase to baseline) were discarded.

#### Proteosome inhibition with MG-132

The fibroblast cells were cultured as normal with 10 µM MG-132 proteasome inhibitor added to the cell media. For the Western blot analysis, the cells were incubated with MG-132 for 12 hours or 24 hours and collected.

#### Quantification and statistical analysis

The statistical analysis was performed with GraphPad Prism 9 (GraphPad Software). The statistical significances between the affected and control cell lines were analyzed with an unpaired two-tailed t-test. The number of replicates is indicated in the figure legends. Significance is defined as *p < 0.05, **p < 0.01, ***p < 0.001 and ****p < 0.0001 in all graphs.

## ACKNOWLEDGEMENTS

Sini Karjalainen, Jana Pennonen and Miia Nissilä (at the University of Helsinki), Anja Florizoone, and Tomas Luyten (at KU Leuven) are acknowledged for technical assistance. We acknowledge the Institute for Molecular Medicine Finland core facility (FIMM) at the University of Helsinki for sequencing services and the IT Center for Science Ltd. (CSC, Finland) for providing a high-performance computing infrastructure.

## Supplementary material

**Supplementary figure 1.**
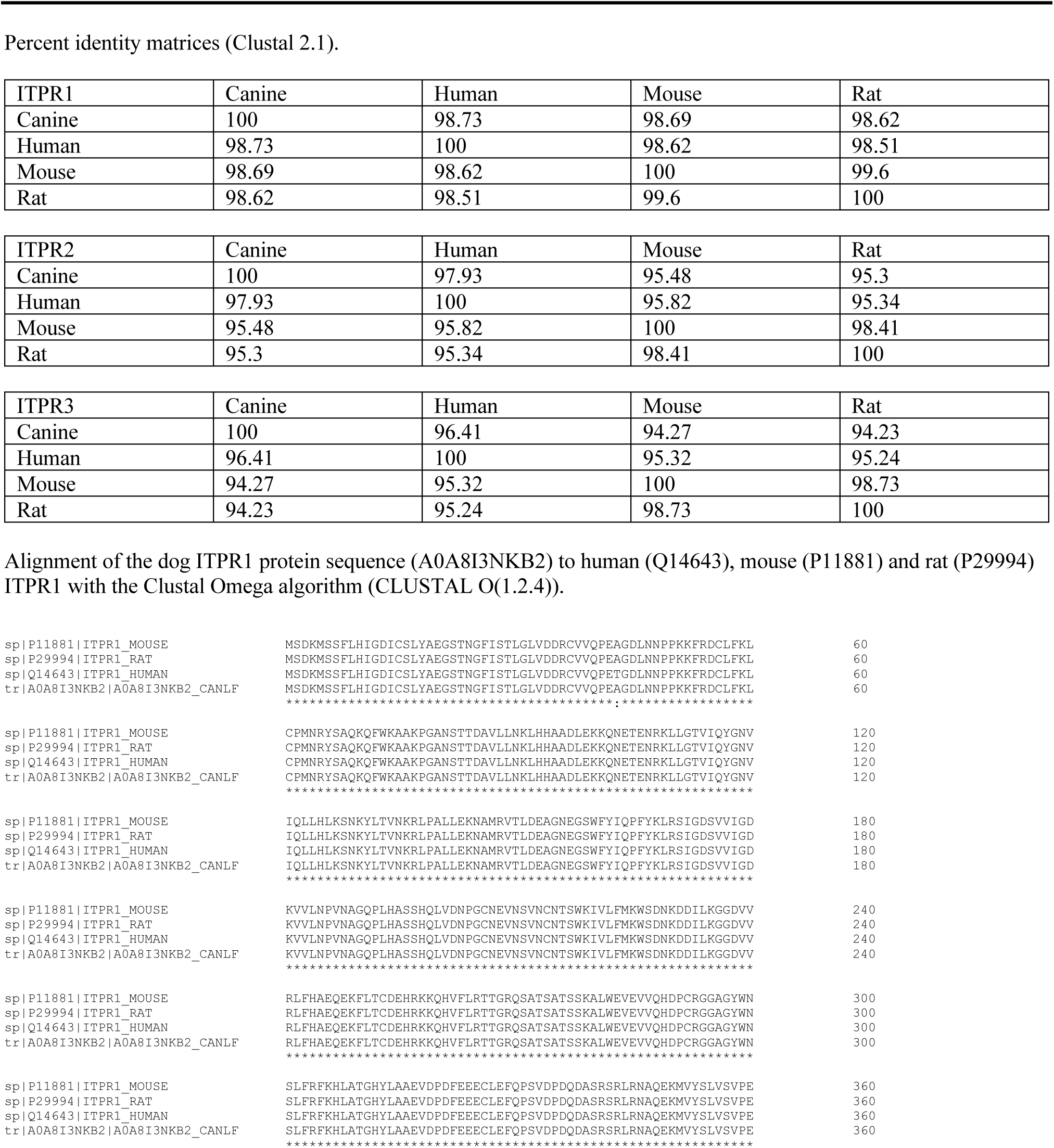
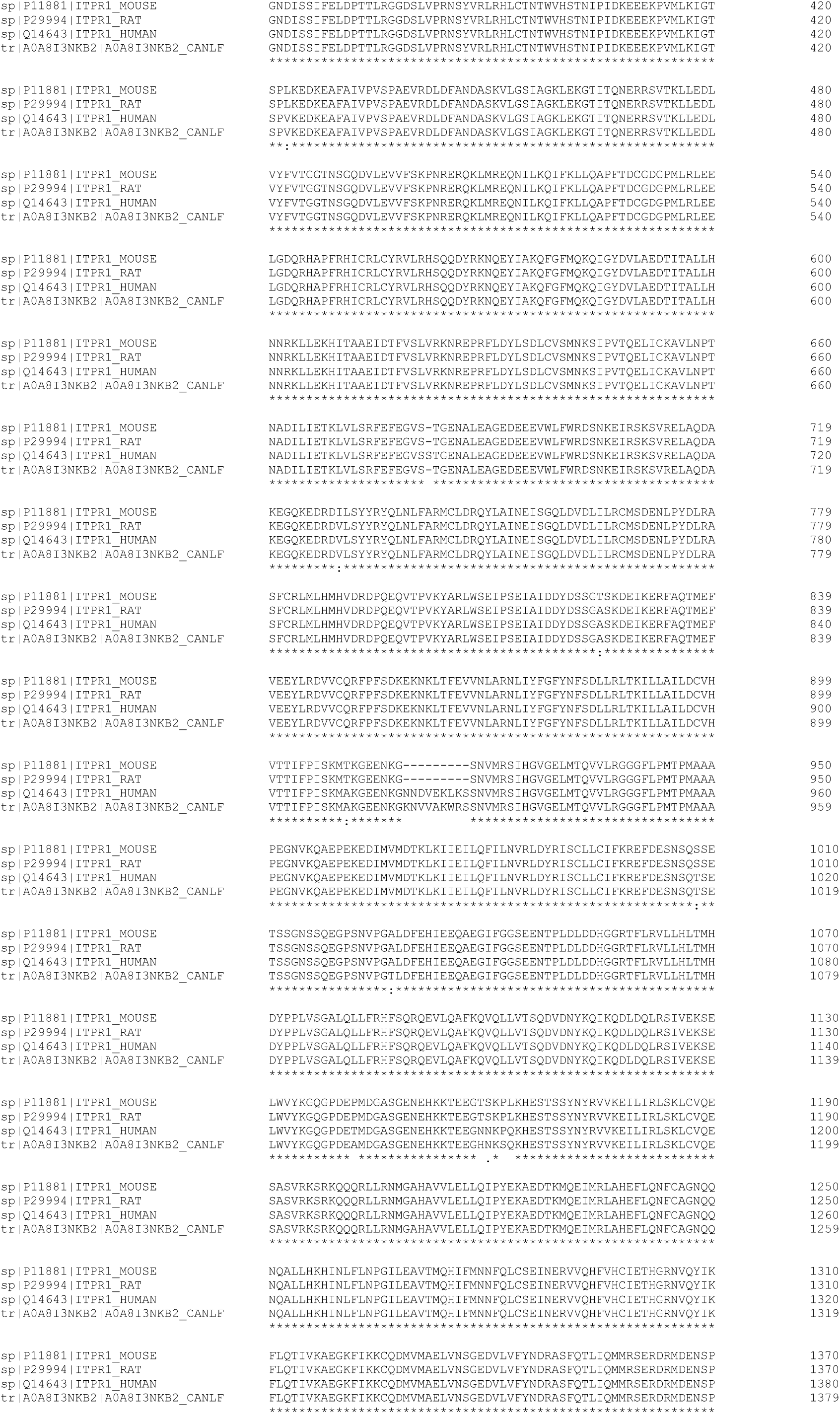
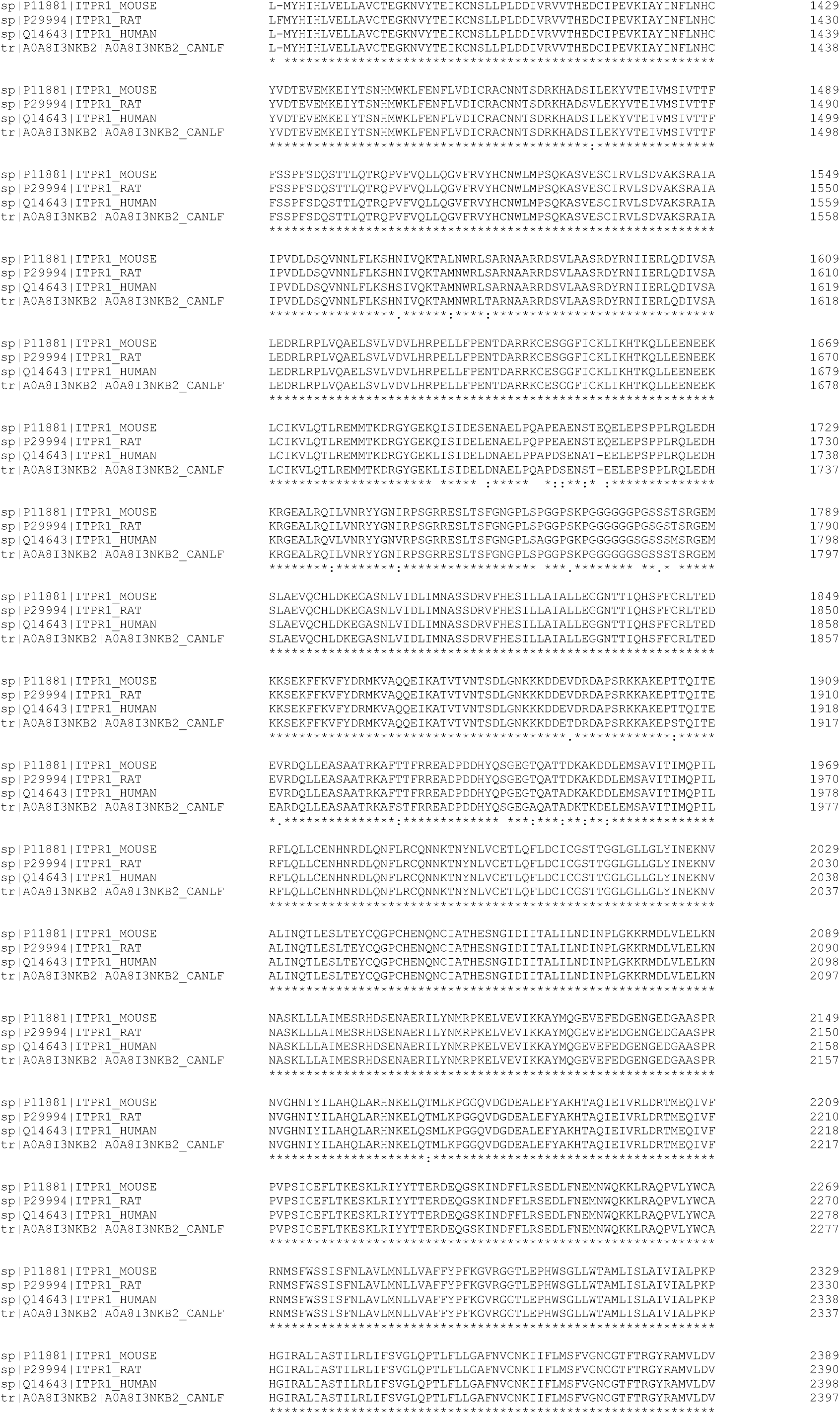
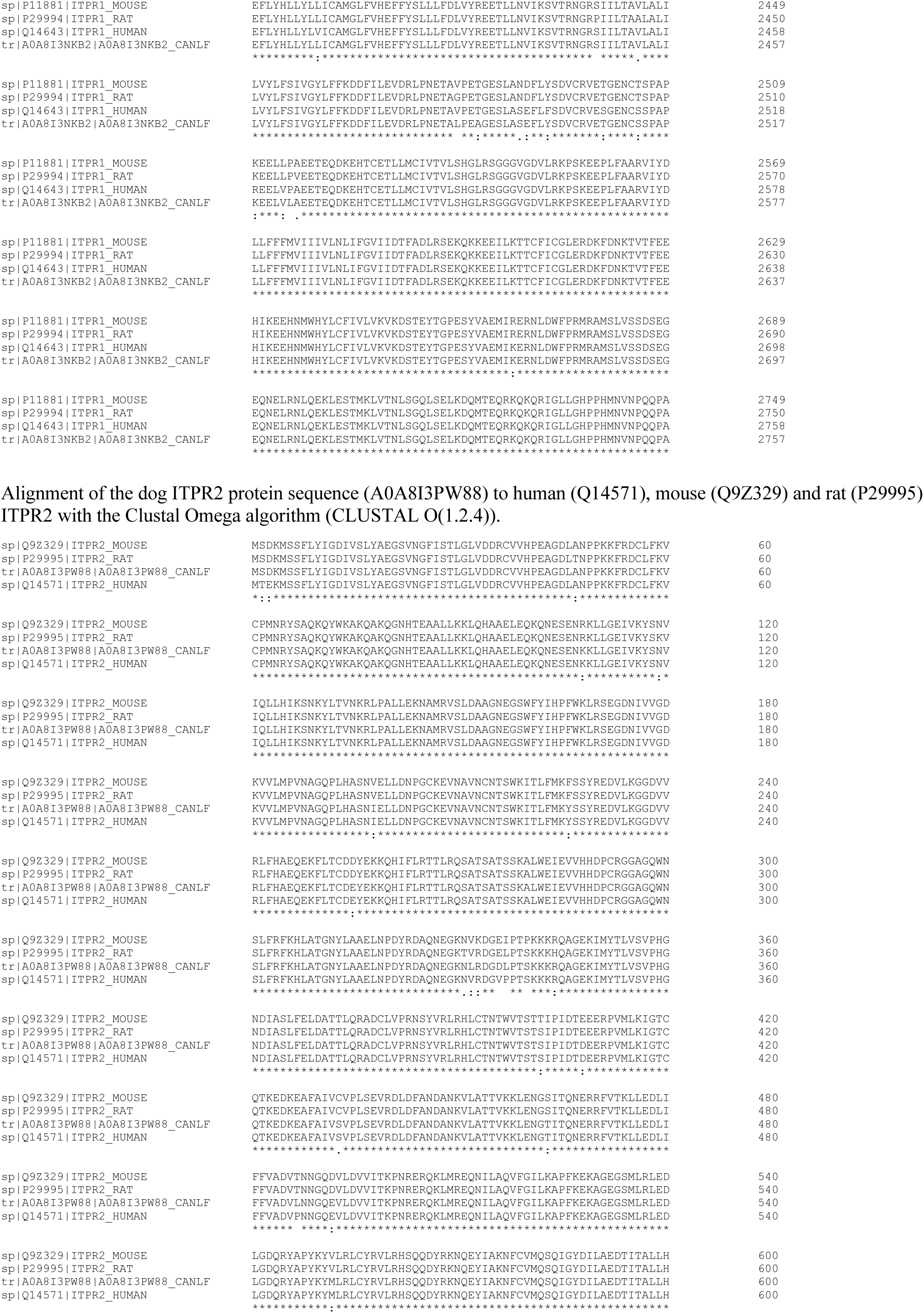
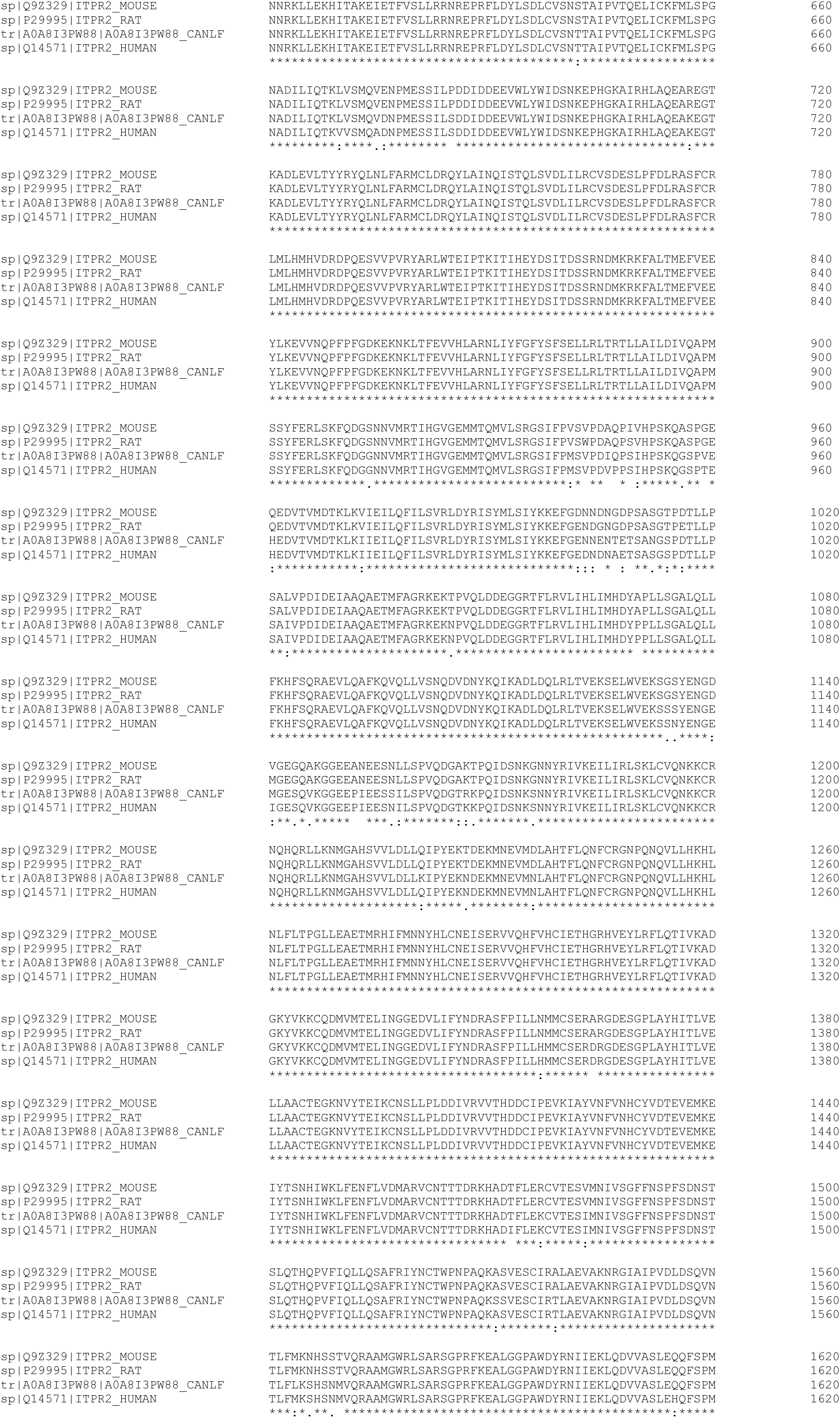
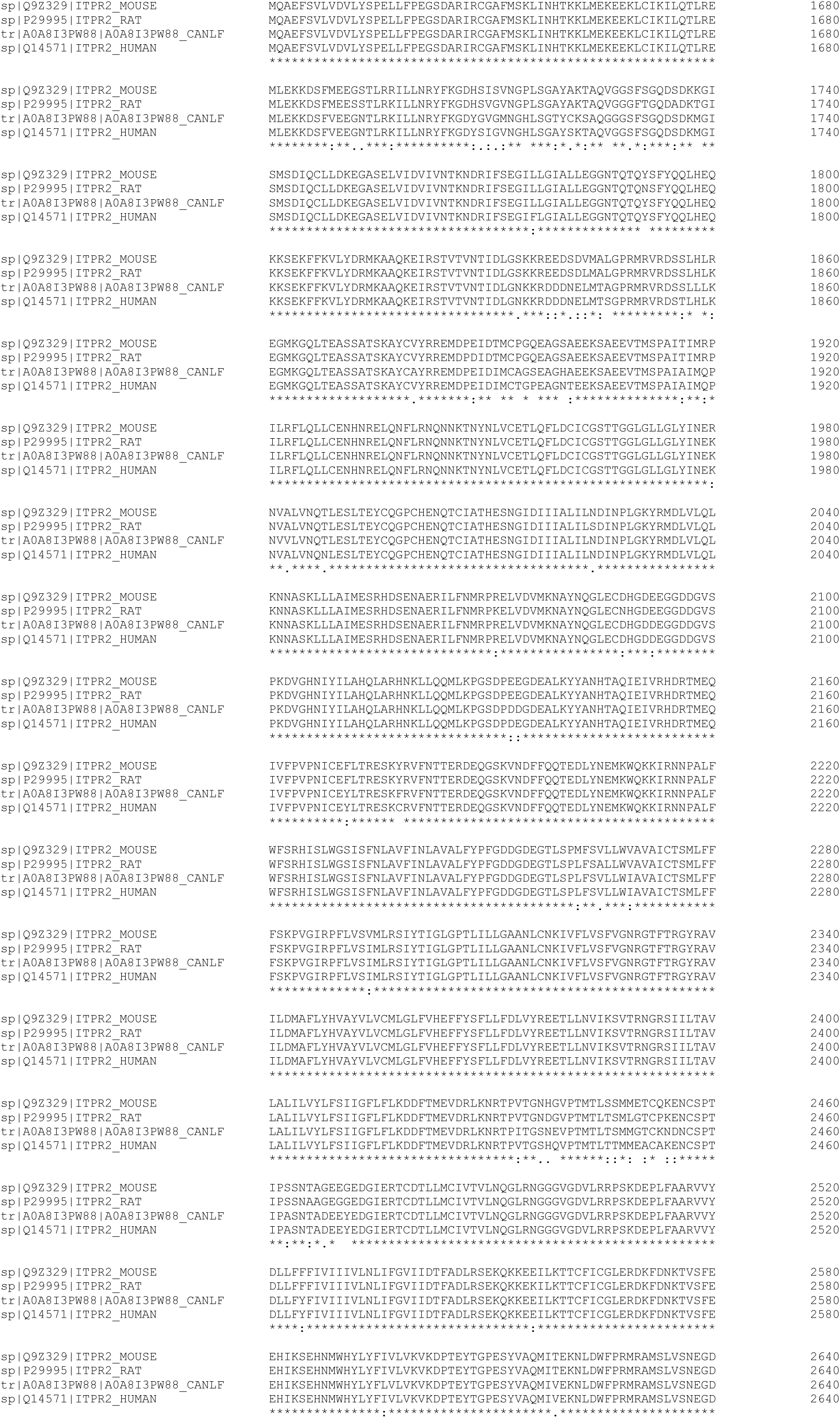
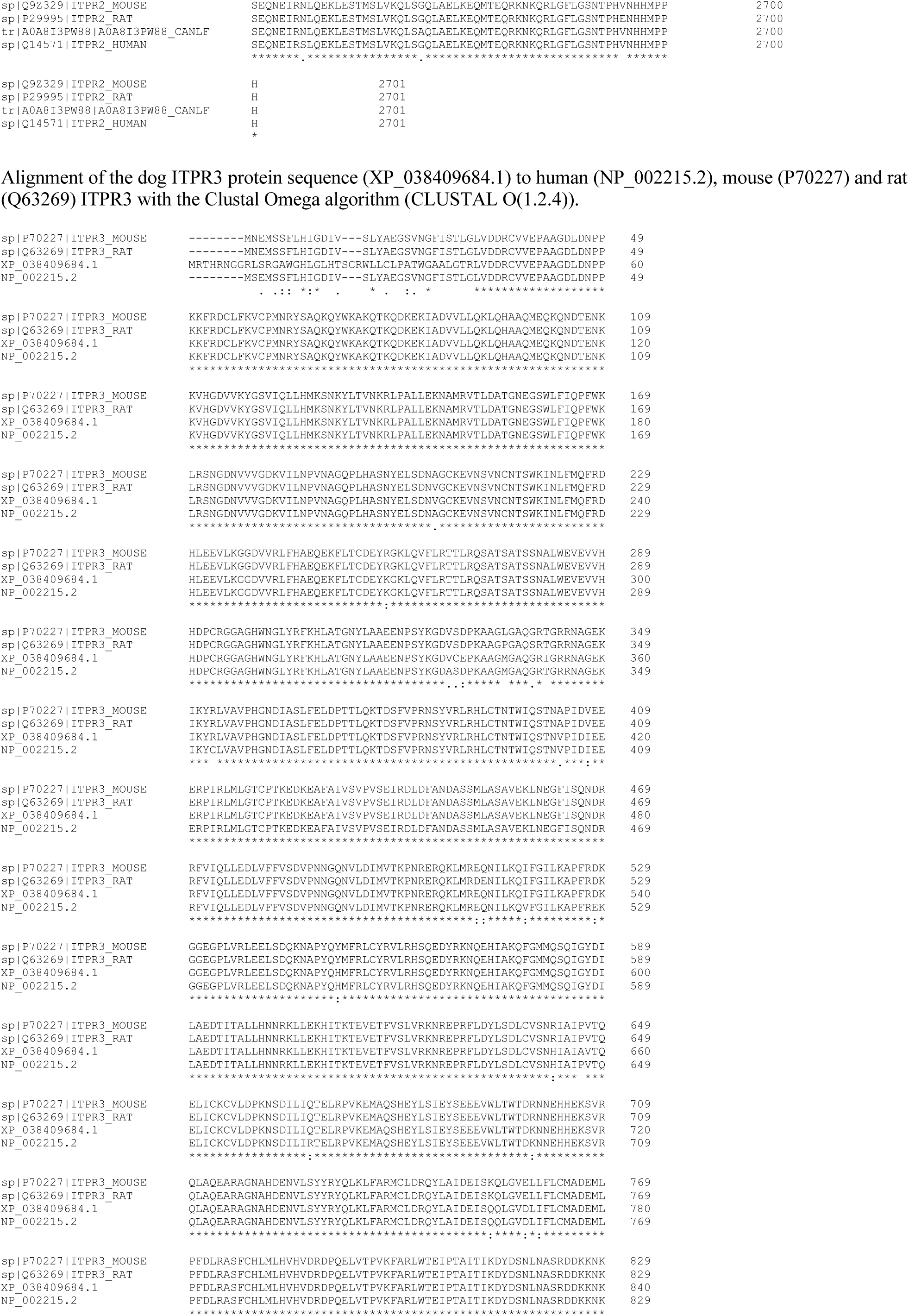
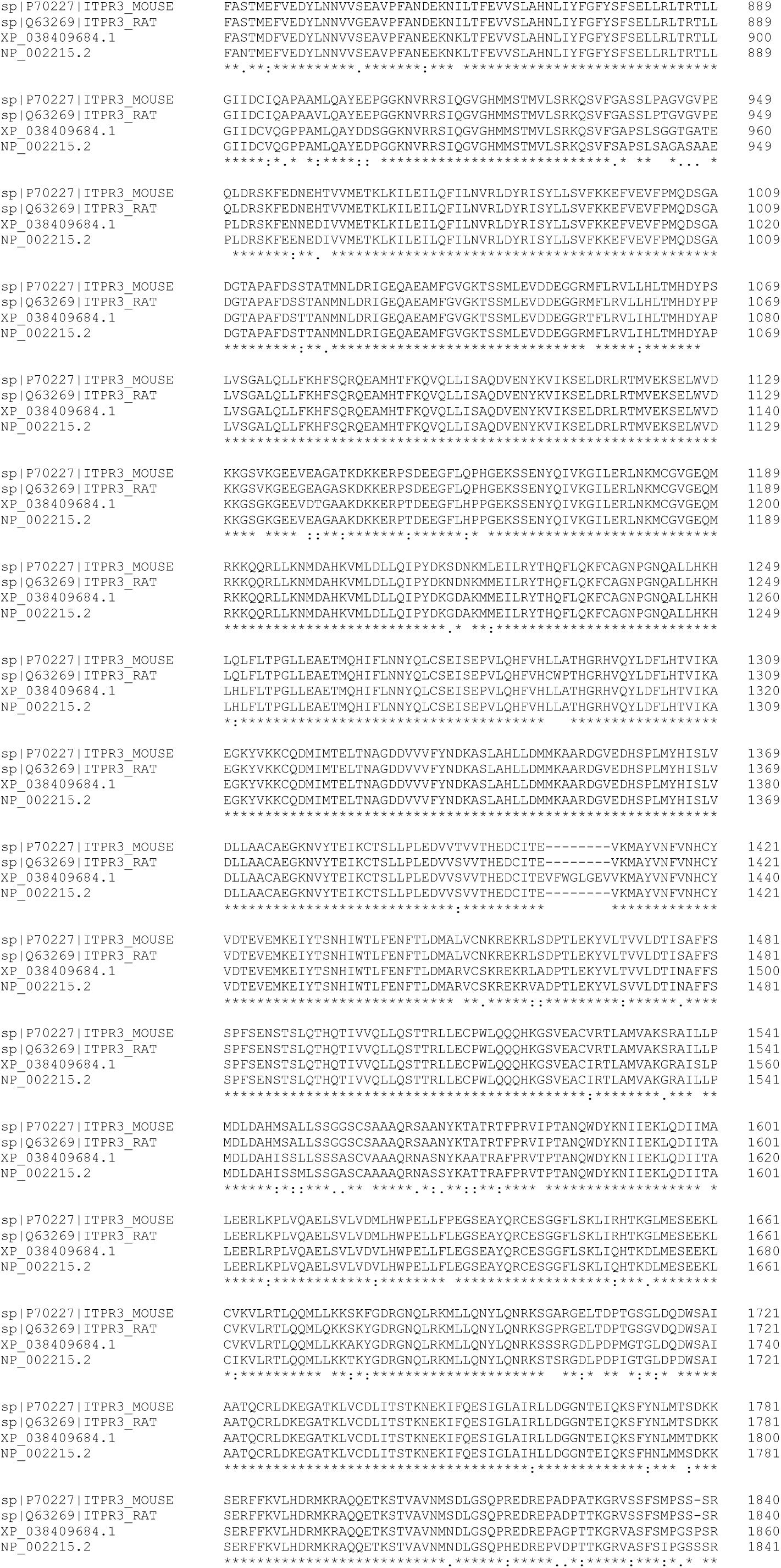
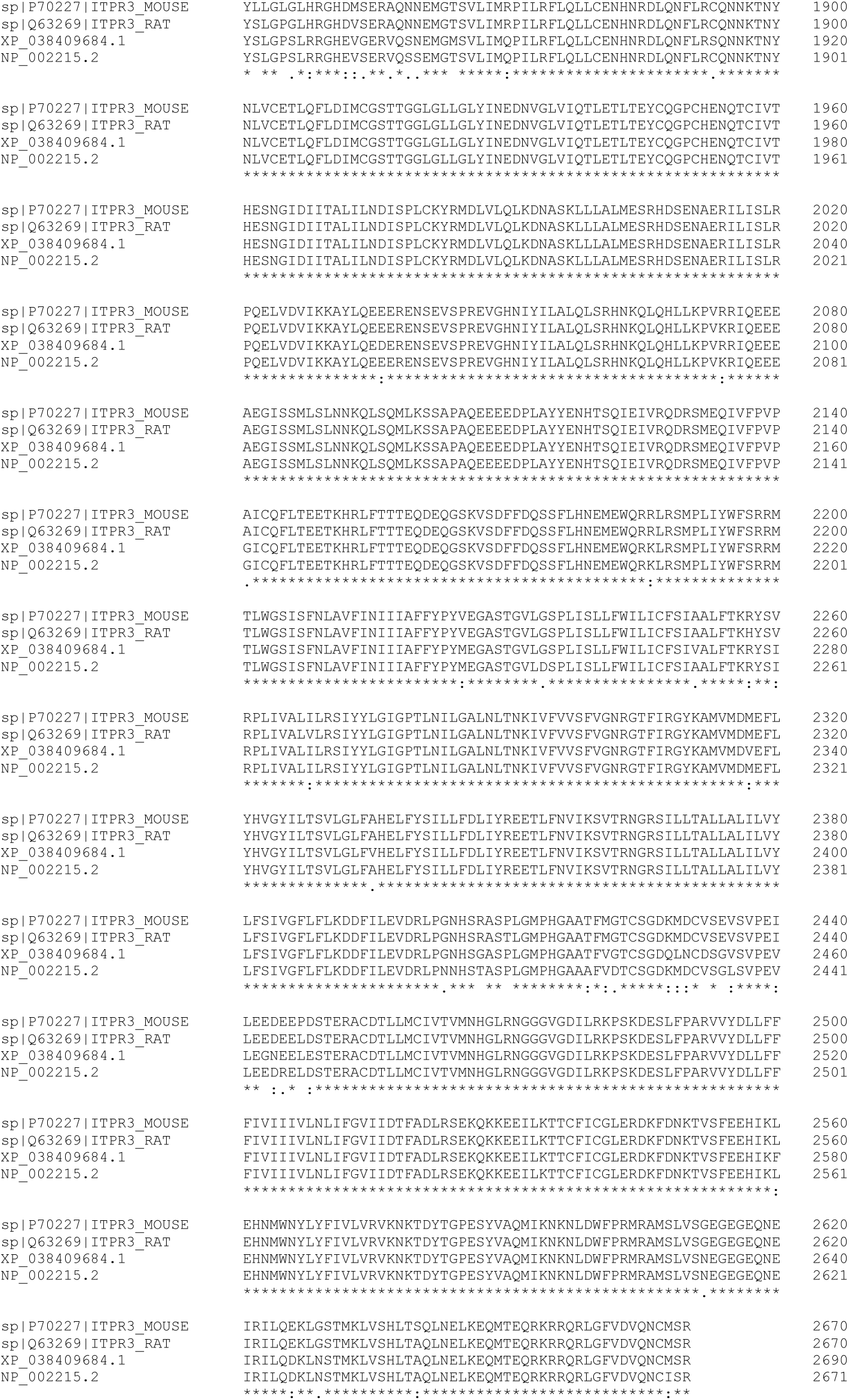
Alignment of the dog IP_3_R1-3 protein sequences to human, mouse and rat IP_3_R1-3 sequences. Nonsense variant in ITPR3 leads to depletion of IP3 receptors in dogs with developmental dental defect and decreased nerve conduction velocity

**Supplementary figure 2.**
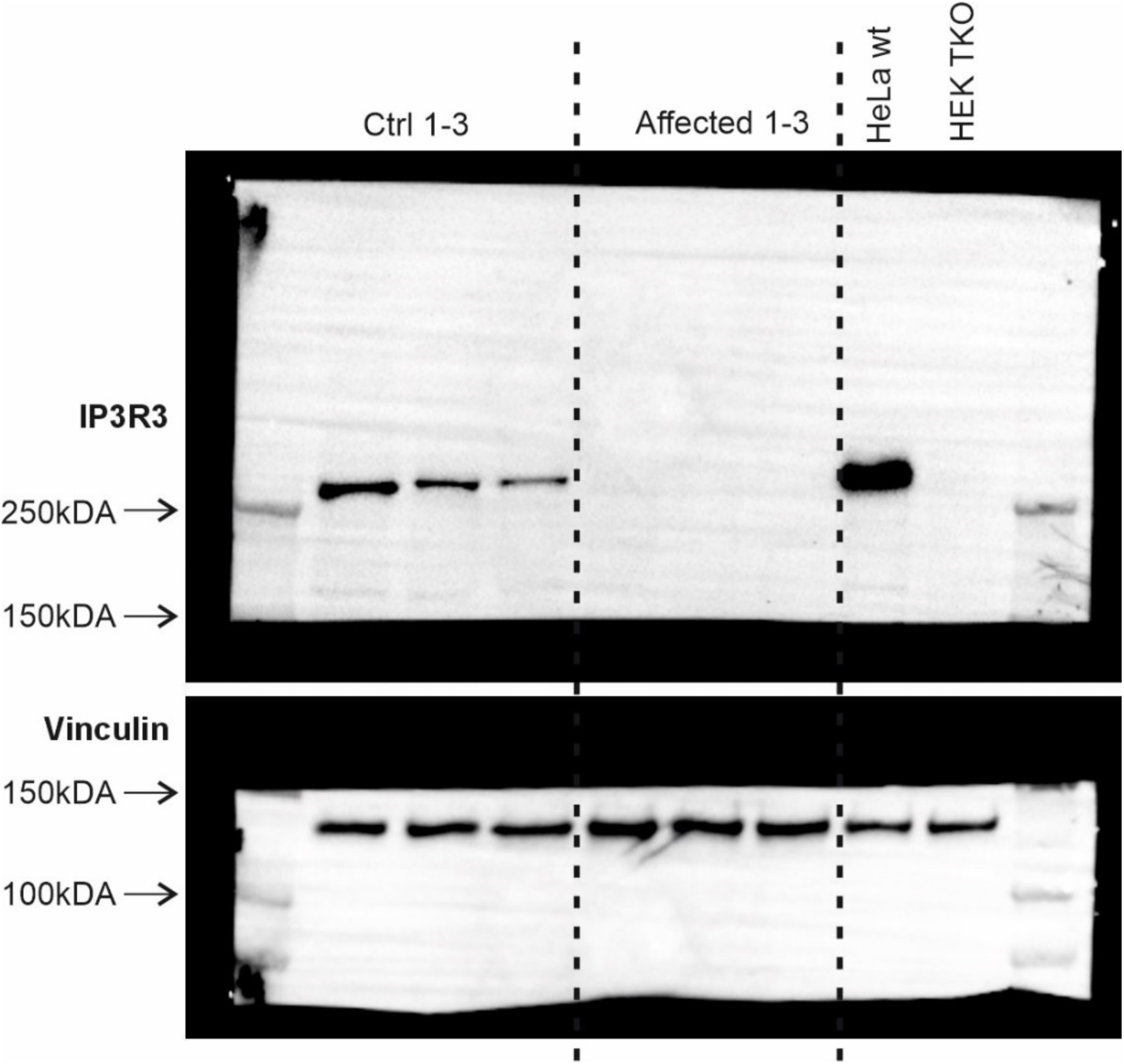
A Western blot of IP_3_R3 protein. The potential truncated IP3R3 protein with the nonsense mutation at p.Q1668, would correspond to a protein with 1667 amino acids if translated. The theoretical size of the protein would be 170-185 kDA. In the Western blot, there are no bands visible on the affected dogs’ fibroblasts at the region corresponding to 150-300 kDA. Lanes: 1-3: healthy control dogs’ fibroblasts, 4-6: affected dogs’ fibroblasts, 7: HeLa wildtype, 8: triple-KO (missing IP_3_R1-3) HEK293.

**Supplementary figure 3.**
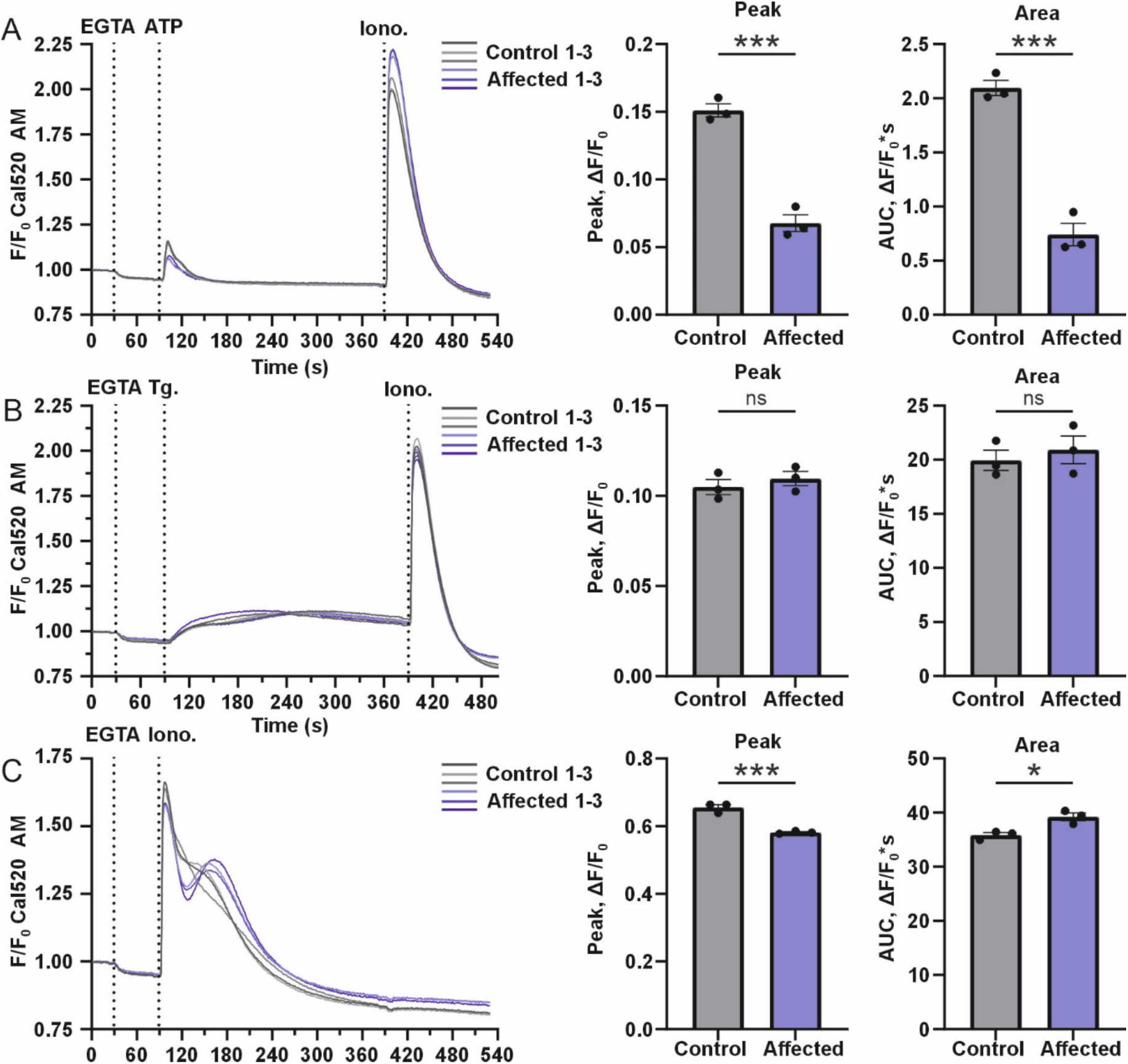
Ca^2+^ flux measurements with drug screening system FDSS/µCell. The Ca^2+^ flux measurements were performed with FDSS/µCell functional drug screening system, using cell permeant Cal-520 AM fluorescent Ca^2+^ indicator. **(A)** Fluorescent response curves of the affected and control cell lines after stimulation with GPCR agonist ATP. The cells were stimulated with 3 mM Ca^2+^ chelator EGTA, 100 µM ATP and 2.5 µM ionomycin. In response to GPCR agonist ATP, the affected fibroblast cells had significantly reduced peak amplitude and area under curve (AUC) (n = 3). **(C)** Fluorescent response curves of the affected and control cell lines after stimulation with 3 mM EGTA, 2 µM thapsigargin and 2.5 µM ionomycin. The loading of ER Ca^2+^ storages were measured with SERCA inhibitor thapsigargin. There were no significant differences in the peak amplitude and AUC between the affected and control cell lines (n = 3). **(D)** Fluorescent response curves of the affected and control cell lines after stimulation with 3 mM EGTA and 2.5 µM ionomycin. The total intracellular Ca^2+^ levels were measured with Ca^2+^ ionophore ionomycin. The affected cells had reduced peak amplitude, while the AUC was elevated in comparison to control cells (n = 3). Data are presented as mean ± SEM. Statistics are analyzed by Student’s t-test (*p < 0.05, **p < 0.01, ***p < 0.001).

**Supplementary table 1.** Electromyographic findings of the affected dogs.

| Lancashire Heeler | Ulnar<br>(normal >60 m/s) | Ishiadicus/peroneal<br>(normal >80 m/s) | Fibrillation<br>potentials | Positive<br>sharp waves | CMAP<br>amplitude | CMAP<br>duration |
| --- | --- | --- | --- | --- | --- | --- |
| Affected 1<br>(age 10y 5m) | 58 m/s | 62 m/s | + | + | ↓ | ↑ |
| Affected 2<br>(6y 10m) | 44 m/s | 76 m/s | + | + | ↓ | ↑ |
| Affected 3<br>(6y 5m & 7y 6m) | 80 m/s | 95 m/s | + | + | - | - |

**Supplementary table 2.** Private variants (SNVs/indels) of three affected dogs after filtering the data against 689 control genomes.

**Supplementary table 3.** Private variants (SNVs/indels) of one affected dog genome after filtering it against 689 control genomes.

**Supplementary table 4.** Private mobile element insertions (MEI) of one affected dog genome after filtering it against 388 control genomes.

**Supplementary tables 2-4.**

Supplementary tables 2-4 are in .xlsx format (Excel sheets).

**Supplementary table 5.** Values used for parameter optimization and detection of shared ROH in PLINK.

|  | values by analysis type |  |
| --- | --- | --- |
| parameter | parameter optimization | detection of shared ROH |
| --homozyg-window-snp | 20 | 20 |
| --homozyg-window-missing | 1 | 1 |
| --homozyg-window-het | 0 | 1 |
| --homozyg-window-threshold | 0.05 | 0.05 |
| --homozyg-snp | 70 | 54 |
| --homozyg-kb | 1000 kb | 1000 kb |
| --homozyg-gap | 2000 kb (when unvaried) | 200 kb |
|  | 20-1000 kb (when varied) |  |
| --homozyg-density | 200 kb/snp (when unvaried) | 25 kb/snp |
|  | 10-125 kb/snp (when varied) |  |

**Supplementary table 6.**
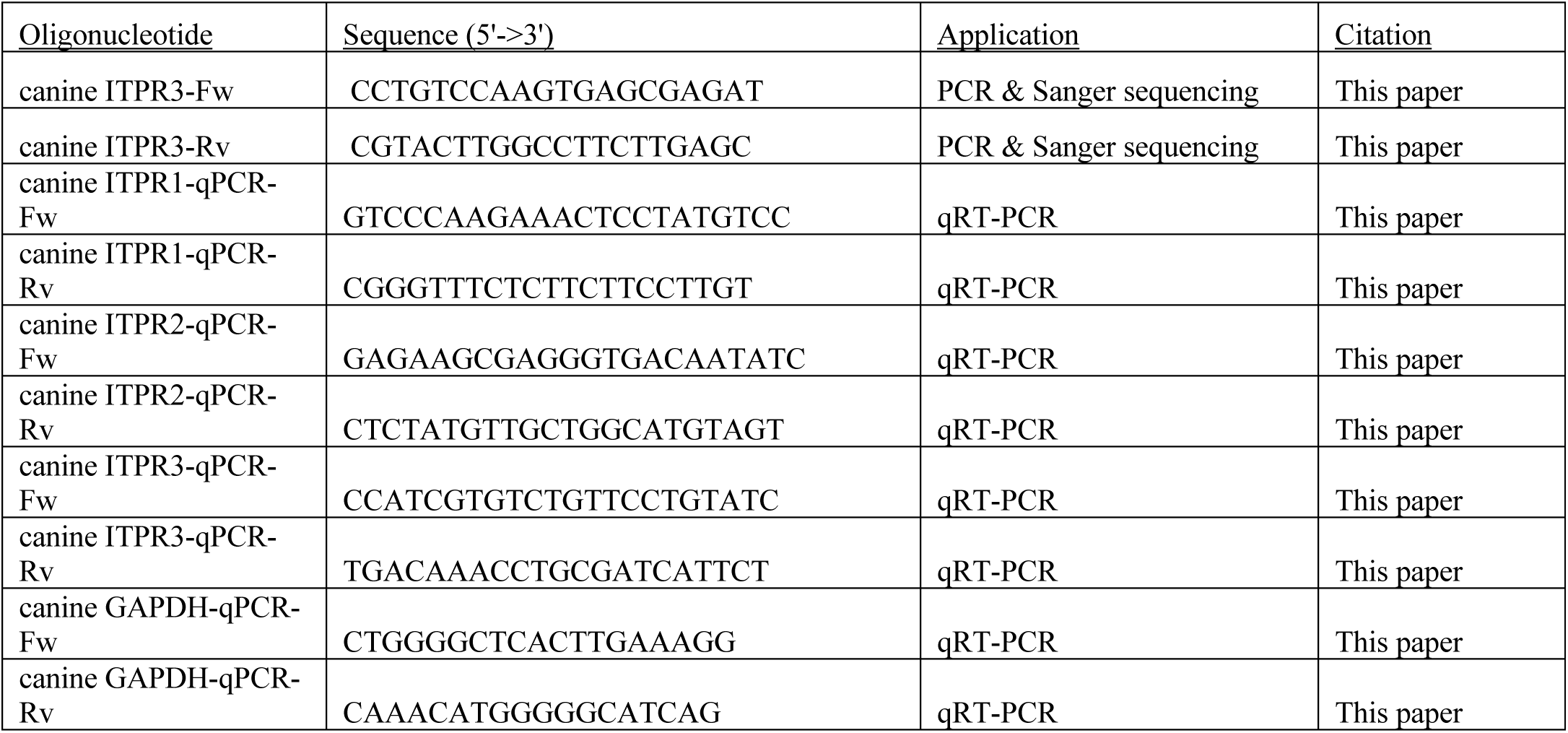
Primers used in the study.

